# Off-target effects of PLX5622 revealed: mixing microglial function in anesthesia and addiction withdrawal

**DOI:** 10.1101/2025.07.18.665493

**Authors:** Kelei Cao, Wang Cheng, Liyao Qiu, Zexi Wang, Yuqing Zhao, Ying Yuan, Weiying Wu, Jingyuan Xue, Linghui Zeng, Zhi-Ying Wu, Huan Ma, Tingjun Hou, Cunqi Ye, Zhihua Gao

## Abstract

Microglia have gained increasing attention as important regulators in the brain. Acute microglial depletion using the colony-stimulating factor 1 receptor (CSF1R) inhibitor, PLX5622, has become a popular approach to uncover microglial function. PLX5622 treatment has been recently shown to significantly promote anesthetic emergence. In characterizing underlying mechanisms, we found that PLX5622 treatment did not change the global neuronal activity during anesthesia, but robustly enhanced anesthetic metabolism. Moreover, PLX5622 dramatically induced the expression of hepatic enzymes that extensively modify xenobiotic and endobiotic metabolism. Blocking increased enzymatic activity significantly reduced the arousal-promoting effects of PLX5622 in anesthesia, irrespective of microglial elimination. In addition, we demonstrate that enhanced drug metabolism also contributed to PLX5622-induced attenuation in nicotine-withdrawal anxiety. By revealing previously unrecognized effects of PLX5622, our findings raise caution in interpreting data generated from PLX5622 treatment and bring forward the need for designing more specific CSF1R inhibitors.

## Introduction

Microglia, the immune sentinels in the central nervous system (CNS), continuously survey the microenvironment with their motile processes to maintain brain homeostasis ^1^. With the advance in microglial manipulation tools, accumulating studies have uncovered diverse functions of microglia, including both immune and non-immune roles, in regulating brain physiology and pathology ^2–8^. Among these tools, pharmacologic blockade of the colony-stimulating factor 1 receptor (CSF1R), a key molecule for microglial survival, inducing rapid microglial death with no apparent behavioral and inflammatory abnormalities in mice ^9–11^, has become a frequently-used approach for functional characterization of microglia, tests of microglial transplantation in neurodegenerative diseases, and combinatorial immunotherapy in cancers ^12–17^.

Using these pharmacologic tools, microglia have been shown to exhibit diverse functions in multiple neurological and psychiatric disorders, including seizure, anesthesia, addiction and anxiety ^7, 18–25^. While predominant studies have highlighted a role for microglia in modulating brain functions by intricate interplays between microglia and neurons ^2, 26–28^, recent reports have pointed out potential caveats and side effects of different microglia manipulation tools. For example, PLX5622, the most potent CSF1R inhibitor, may also target oligodendrocyte precursor cells and endothelial cells in the brain, in addition to peripheral macrophages ^29–32^, raising critical concerns whether outcomes observed upon PLX5622 treatment are faithfully reflective of microglial functions.

The roles of microglia in modulating anesthesia and sleep are increasingly gaining attention ^7, 18, 26, 28, 33, 34^. Intriguingly, although cortical microglia exhibit similar changes during NREM/REM sleeps and anesthesia ^27, 35–38^, microglial depletion have distinct consequences on sleep and anesthesia. PLX5622 or another CSF1R inhibitor, PLX3397 treatment does not alter the overt sleep-wake transitions ^26, 33, 34^, but only transiently disturbed sleep within a narrow temporal window ^26, 33^. By contrast, PLX5622 treatment dramatically accelerated anesthetic emergence from different types of injectable anesthetics, regardless of mechanisms of actions ^39^. How microglia achieve to elicit such differential effects on sleep and anesthesia remains a conundrum.

To further identify mechanisms underlying microglial contributions in anesthesia and sleep, we found that PLX5622 treatment only affected injectable anesthesia, with no apparent effects on inhalation anesthesia and sleep. The selective impact of PLX5622 treatment on injectable anesthetic arousal is primarily attributable to the robust upregulation of metabolic enzymes in the liver, which drives anesthetic metabolism. Such hepatic responses substantially enhanced metabolism of xenobiotics and also contributed to the anxiety attenuation upon nicotine withdrawal after PLX5622 treatment. Given that CSF1R inhibitors have been extensively applied to investigate microglia and macrophage-related scientific questions and disease mechanisms, the neglected confounding effects of PLX5622 raise important awareness for rigorous experimental design and accurate data interpretation during its application.

## Results

### PLX5622 treatment does not affect inhalation anesthesia and vigilance states

We and others have recently shown that PLX5622 treatment in mice robustly promoted anesthetic arousal induced by a variety of injectable anesthetics, regardless of their different mechanisms of actions ^7, 18^. Since PLX5622 treatment induces microglial elimination, these effects were attributed to microglial regulation. In testing the roles of microglia in inhalation anesthesia **(Fig. 1a)**, however, we observed no differences in the induction and recovery time of both isoflurane and sevoflurane anesthesia after PLX5622 treatment (in WT mice) or diphtheria toxin (DT)-treatment (*in Cx3cr1^CreER/^+::iDTR^fl/fl^*mice) **(Fig. 1b-e, Extended Data Fig. 1a-c)**. EEG recordings also showed no difference in the loss and restoration time of consciousness or power of δ and α/β waves during isoflurane anesthesia, between WT and PLX5622-treated mice **(Fig. 1f-j, Extended Data Fig. 1d-f)**. Further assessment by continuous EEG recording revealed that PLX5622 treatment did not change the overall vigilance states and sleep architecture in mice over 24 hr period **(Fig. 1k-q, Extended Data Fig. 2a-r)** ^34^, with only slightly increased NREM/REM sleep (NREM: 31.18±6.08% *v.s.* 49.1±4.41%, *p=0.042*; REM: 2.81±0.84% *v.s.* 6.33±1.11%, *p=0.025*) and reduced wakefulness (66.00±6.91% *v.s.* 44.67±5.36%, *p=0.036*) during the light-off stages of ZT19-21 (**Extended Data Fig. 1g-i**). Together, these data demonstrate that PLX5622 treatment had no significantly impacts on inhalation anesthesia and overall vigilance states, despite of its striking influence on injectable anesthesia.

**Fig 1.**
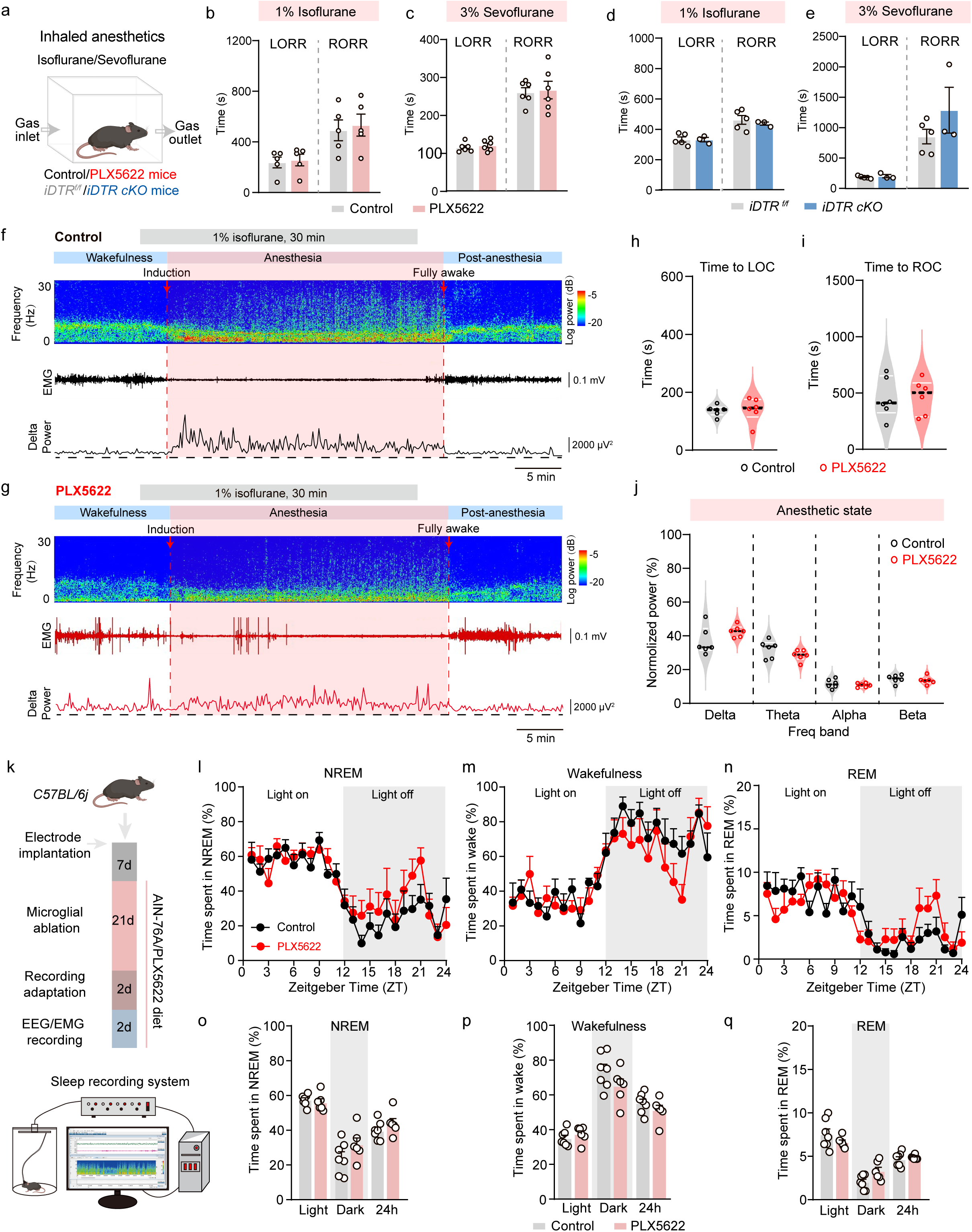
Microglial depletion barely affected inhalation anesthesia and slightly altered the sleep patterns. **(a)** A scheme of experimental design and inhalation anesthesia tests. **(b-c)** Time of anesthetic induction (loss of righting reflex, LORR) and recovery (recovery of righting reflex, RORR) after inhalation of 1% isoflurane (b) (n=5 mice per group), and 3% sevoflurane (c) in control and microglia-depleted (PLX5622-treated) male mice (n=6 mice per group). **(d-e)** Time of LORR and RORR after 1% isoflurane **(d)**, and 3% sevoflurane **(e)** inhalation between control (*iDTR^f/f^*) and microglia-depleted (DT-treated) male mice (n=5 mice for *iDTR^f/f^* and n=3 for *iDTR cKO* group). **(f-g)** Example traces of EEG and EMG recording of control (f) or PLX5622-treated (g) mice after 1% isoflurane exposure. **(h-i)** Time to loss of consciousness (LOC) **(h)** and recovery of consciousness (ROC) **(i)** in control and PLX5622-treated mice (n=6 mice per group). **(j)** Power distribution of EEG frequency bands during 1% isoflurane-induced anesthesia in control and PLX5622 mice (n=6 mice per group). **(k)** A scheme of experimental design and sleep recording paradigm. **(i-n)** Circadian variation of NREM (i), wakefulness (m), and REM (n) in the control and PLX5622 group. **(o-q)** Percentages of NREM (o), wakefulness (p), and REM (q) during the overall 24 h recording course in control and PLX5622-treated mice. n=7 mice for control and n=6 mice for PLX5622 group. Data are represented as mean ± SEM (bar graphs) and median ± interquartile range (violin plots). Unpaired Student’s *t* test for **(b-e, h-j, o-q)** \**p* < 0.05, \*\**p* < 0.01, and \*\*\**p* < 0.001. NREM, non-rapid eye movement, REM, rapid eye movement.

### PLX5622 treatment does not alter neuronal activities and extracellular adenosines during anesthesia

To examine the potential mechanisms underlying the PLX5622-induced anesthetic differences, we performed c-fos screening in the anesthetized brain. However, in both dexmedetomidine or pentobarbital-induced anesthesia, we were unable to detect significant differences in the number of c-fos labeled neurons between WT and PLX5622-treated samples **(Extended Data Fig. 3a-e and data not shown)**, including anesthesia-activated brain regions, such as the amygdala and bed nucleus of stria terminalis ^40–42^, regardless of microglial presence.

An important mechanism for microglia to dampen neuronal activity is adenosine production ^2^. To test whether PLX5622 treatment changes the dynamics of extracellular adenosine in the anesthetized brain as result of microglia elimination, we injected the genetically encoded adenosine sensor (GRAB_Ado_) into the medial prefrontal cortex, a region critical for the regulation of consciousness and cognition ^43^, and tracked the dynamics **(Extended Data Fig. 4a-c**). While both inhalation (isoflurane, **Extended Data Fig. 4d-e**) and injectable (ketamine, **Extended Data Fig. 4i-m**) anesthesia induced robust elevation of extracellular adenosine, PLX5622 treatment did not significantly change the amount of adenosine during steady-state anesthesia (0.43±0.07 *v.s.* 0.58 ± 0.07 for isoflurane, *p=0.1173*; 0.34 ± 0.06 *v.s.* 0.33 ± 0.05 for ketamine, *p=0.8707*) **(Extended Data Fig. 4d-s)**. These data suggest that PLX5622 treatment does not induce global changes in neuronal activity and extracellular adenosine in the anesthetized brain.

### PLX5622 treatment promotes drug biotransformation

The distinct effects of PLX5622 treatment on injectable and inhalation anesthesia, with no apparent changes in neuronal activity, prompted us to pursue non-neural mechanisms underlying these interesting phenomena. A critical difference between inhaled and injectable anesthetics lies in their pharmacokinetics: inhaled agents undergo rapid respiratory absorption/elimination, whereas injectable drugs rely on hepatic metabolism and renal excretion **(Fig. 2a)** ^44, 45^. Since inhalation anesthesia involves no hepatic metabolism, we speculated that PLX5622 treatment may alter drug metabolism in the liver to selectively affect injectable anesthesia.

**Fig 2.**
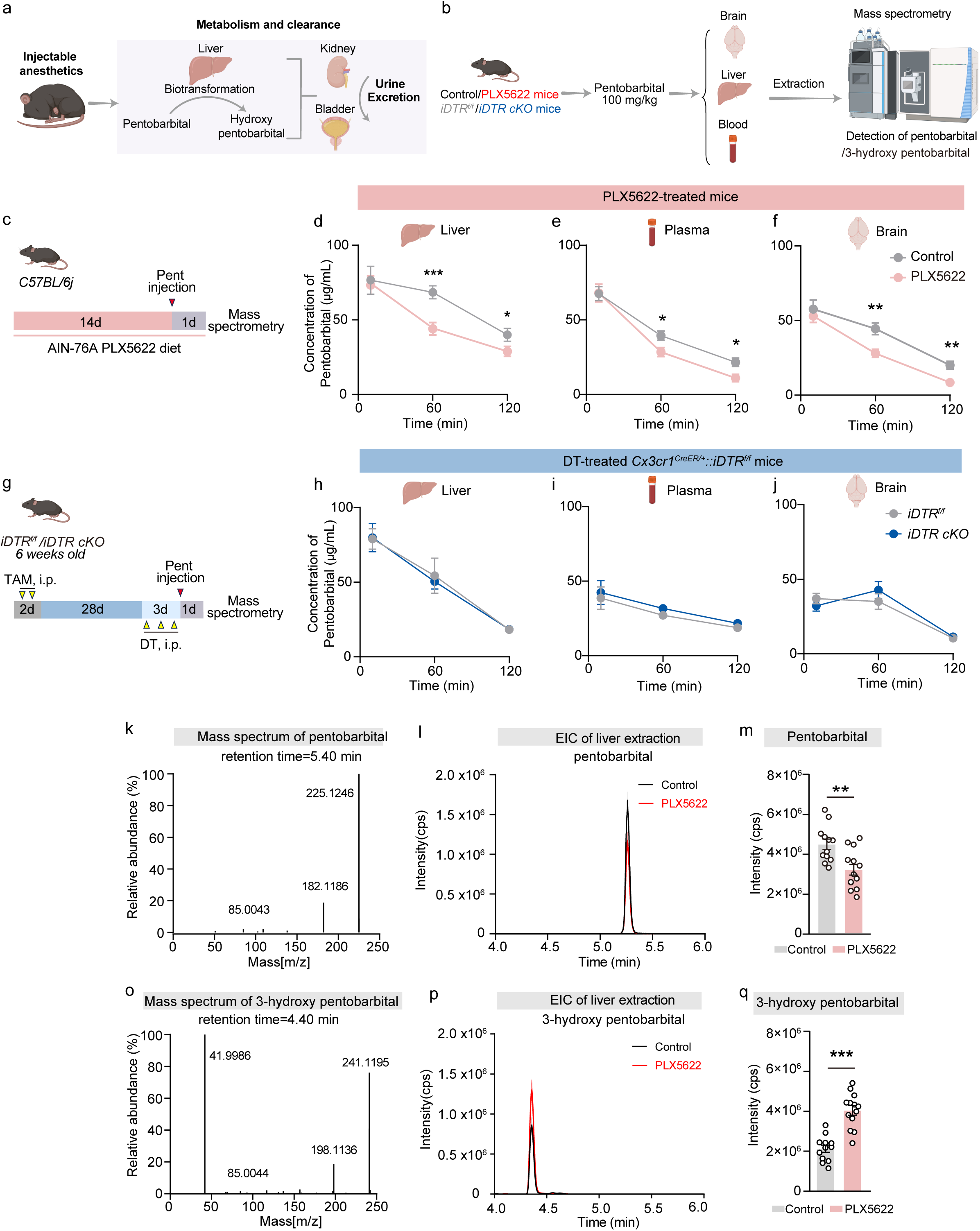
Pharmacologic (PLX5622-treated) depletion of microglia promoted pentobarbital metabolism. **(a)** A scheme of biotransformation, metabolism and clearance of pentobarbital. **(b)** A scheme of microglial depletion and subsequent analyses. **(c)** An experimental timeline. **(d-f)** Pentobarbital concentrations in the liver **(d)**, plasma **(e)**, and brain **(f)** in control and microglia-depleted (PLX5622-treated) mice following pentobarbital injection. n=5-13 for control mice, and n=6-9 for PLX5622 mice. **(g)** The experimental timeline and timing of tamoxifen administration, diphtheria toxin (DT) administration, pentobarbital injection, and mass spectrometry detection. **(h-j)** Dynamics of pentobarbital concentrations in the liver **(h)**, plasma **(i)**, and brain **(j)** in *iDTR^f/f^* and microglia-depleted (*iDTR cKO*) mice following pentobarbital injection. n=5-14 for *iDTR^f/f^* mice, and n=5-10 for *iDTR cKO* mice. **(k, o)** Electron impact high-resolution mass spectrometry (MS) and MS fragmentation spectra of pentobarbital (k) and 3-hydroxy pentobarbital (o, hydroxylated metabolite of pentobarbital). **(l)** Extracted ion chromatogram (EIC) showing a significantly higher peak for pentobarbital at a retention time of ∼5.40 min in the control group compared to the PLX5622 group. **(p)** EIC displaying a notably lower peak for 3-hydroxy pentobarbital at a retention time of ∼4.40 min in the control group compared to the PLX5622 group. **(m, q)** The particle counts of pentobarbital **(m)** and its hydroxylated metabolite, 3-hydroxypentobarbital **(q)** in the liver of control and microglia-depleted (PLX5622-treated) mice at 1 h after pentobarbital injection. n=12 mice for controls, n=13 mice for PLX5622-treated group. Data are represented as mean ± SEM (bar graphs). Unpaired Student’s *t* test for (**d-j, m-q**), \**p* < 0.05, \*\**p* < 0.01, and \*\*\**p* < 0.001. Pen: pentobarbital; TAM: tamoxifen; DT: diphtheria toxin.

To test this possibility, we measured the concentration of pentobarbital in the livers, plasma and brains of WT and PLX5622-treated mice at different time points after drug administration **(Fig. 2b)**. At 10 min post-injection, a time point with peak accumulation of pentobarbital ^46^, we observed no differences in the amounts of pentobarbital in between controls and PLX5622-treated mice **(Fig. 2c-f)**. By 1-2 h, however, there is a remarkable decrease of pentobarbital in the liver (1 h: 68.48±4.37 *v.s.* 44.05±4.16 μg/ml, *p=0.0008;* 2 h: 40.07±10.75 *v.s.* 28.91±7.59 μg/ml, *p=0.0116*); plasma (1 h: 39.43±3.07 *v.s.* 28.47±3.01 μg/ml, *p=0.0418*; 2 h: 21.60±2.89 *v.s.* 11.02±2.56 μg/ml, *p=0.0136*); and brain (1 h: 44.44±3.97 *v.s.* 27.96±2.79 μg/ml, *p=0.0044*; 2 h: 20.02 ± 2.60 *v.s.* 8.43±2.00 μg/ml, *p=0.0025*) from PLX5622-treated mice **(Fig. 2c-f)**. The decline of pentobarbital [Control: (4.46 ± 0.26)×10^6^ *v.s.* PLX5622: (3.18 ± 0.27) ×10^6^ counts per second (cps), *p=0.0027*)] was accompanied by a simultaneous increase of its metabolite, 3-hydroxy pentobarbital in the liver [Control: (2.13±0.18)×10^6^ *v.s.* PLX5622: (4.05 ± 0.24) ×10^6^ cps, *p<0.0001*)] at 1 h **(Fig. 2k-q).** By contrast, concentrations of pentobarbital in both liver, plasma and brain remains unchanged between controls and DT-treated *Cx3Cr1^CreER/+^::iDTR^fl/fl^* mice **(Fig. 2g-j)**. Together, these data indicate that PLX5622, but not DT treatment, substantially enhanced hepatic metabolism of pentobarbital.

### PLX5622 induces CYP2B10 expression to drive drug metabolism and reduce anesthetic efficacy

Since PLX5622 treatment promotes pentobarbital biotransformation, and hepatic cytochrome P450 (CYP) enzymes play pivotal roles in xenobiotic biotransformation ^47, 48^, we reasoned that PLX5622 may have induced hepatic cytochrome P450 (CYP) enzymes to elicit such consequences. RT-qPCR analyses of three major anesthetics-metabolizing enzymes (CYP2B10, 2C37 and 3A11) revealed robust hepatic induction ^49–52^, with *Cyp2b10* induced by more than 7000-fold, after two-week treatment of PLX5622 **(Fig. 3a-b)**. By contrast, DT treatment resulted no apparent changes in these enzymes in *Cx3cr1^CreER/+^::iDTR^fl/fl^* mice **(Fig. 3c-d)**. Western blotting also confirmed the selective induction of CYPs in PLX5622-treated, but not DT-treated, mice **(Extended Data Fig. 5a-f)**. These data suggest that CYP induction arises directly from PLX5622 treatment, irrelevant to microglial depletion. Indeed, a careful re-assessment and comparison noticed that PLX5622 had much stronger effects than DT-treatment in promoting emergence from injectable anesthesia **(Fig. 3e)**. To test whether PLX5622 treatment could promote anesthetic emergence in a microglia-dependent manner, we further fed DT-treated *Cx3Cr1^CreER/+^::iDTR^fl/f^*mice with control or PLX5622 chow for only two days **(Fig. 3f-h)**. Consistent with our previous observation ^7^, DT-treated *Cx3Cr1^CreER/+^::iDTR^fl/f^* mice exhibited shortened duration of pentobarbital anesthesia (**Fig 3j-k**, RORR time: 484.2±14.15 min in controls v.s. 394.9±17.23 min in DT-treated mice), but with much weaker effects than PLX5622 treatment (**Fig 3k**). By contrast, PLX5622 treatment further shortened the duration of pentobarbital anesthesia by 235.8 min (approximately 59.72% reduction, RORR time: 394.9±17.23 min in DT-treated mice *v.s.* 159.1±25.65 min in DT+PLX mice) (**Fig. 3j-k**), even in the absence of microglia. In addition, acute feeding of PLX5622 chow for 24 h, causing no apparent loss of microglia (**Fig. 3l, Extended Data Fig. 5i-j**), also promoted anesthetic emergence (**Fig. 3n-p**), suggesting that PLX5622 treatment alone can accelerate anesthesia in a microglia-independent manner.

**Fig 3.**
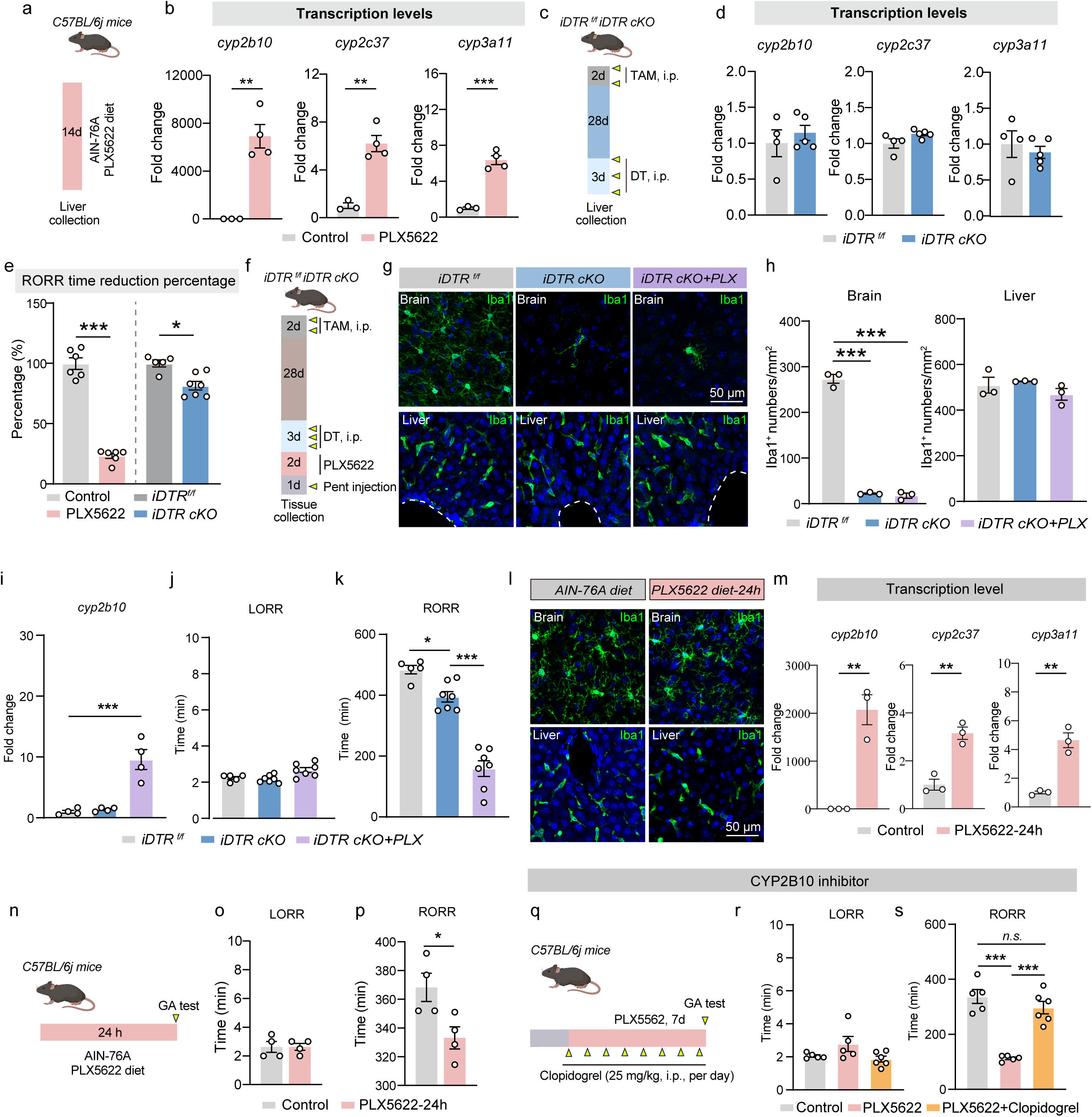
PLX5622 induces CYP2B10 expression to drive drug metabolism and reduce the efficacy of anesthetics. **(a)** An experimental scheme. **(b)** RT-qRCR analysis of the transcription levels of *cyp2b10*, *cyp2c37*, and *cyp3a11* in the liver from control and PLX5622-treated mice. n=3 for control and n=4 for PLX5622-treatment mice. **(c)** An experimental timeline. **(d)** RT-qRCR analysis of *cyp2b10*, *cyp2c37*, and *cyp3a11* in the liver from *iDTR^f/f^* and *iDTR cKO* mice. n=4 mice for *iDTR^f/f^* group, and n=5 mice for *iDTR cKO* group. **(e)** The percentage reduction in pentobarbital anesthetic duration (RORR) in mice with pharmacological or genetic microglial ablation. n=6 mice in control and PLX5622 group, n=5 mice for *iDTR^f/f^* group, n=7 mice for *iDTR cKO* group. (Pharmacological microglial ablation data from our previous paper (*>doi:10.1016/j.cub.2023.04.047 (2023)*, and genetic microglial ablation data from this paper). **(f)** An experimental scheme. **(g)** Representative images of Iba1 staining from the brain and liver of *iDTR^f/f^*, *iDTR-cKO* mice fed with regular or PLX5622 diet as indicated. **(h)** Quantification of microglial number in the cerebral cortex from *iDTR^f/f^*, *iDTR cKO*, and *iDTR cKO* with PLX5622-treated mice. n=3 mice per group. **(i)** RT-qRCR analysis of *cyp2b10* in the liver from *iDTR^f/f^*, *iDTR cKO* mice fed with regular or PLX5622 diet as indicated. n=4 mice per group. **(j-k)** Time of LORR and RORR after intraperitoneal (i.p.) injection of pentobarbital into *iDTR^f/f^*, *iDTR cKO*, and *iDTR cKO* mice treated with PLX5622. (n=5 mice for *iDTR^f/f^*group, n=7 mice for groups of *iDTR cKO* and *iDTR cKO*+PLX). **(l)** Representative images of Iba1 staining from the cerebral cortex and liver following PLX5622 treatment for 24 hours. Iba1^+^ (green) cells represent microglia. **(m)** RT-qRCR analysis of *cyp2b10*, *cyp2c37* and *cyp3a11* in the liver from control, and 24h PLX5622-treated mice. n=3 mice per group. **(n)** An experimental timeline. **(o-p)** Time of LORR **(o)** and RORR **(p)** after i.p. injection of pentobarbital (100 mg/kg) between control and 24h PLX5622-treated mice. n=4 mice per group. **(q)** An experimental timeline. **(r-s)** Time of LORR **(r)** and RORR **(s)** after i.p. injection of pentobarbital between control, PLX5622-treated groups with or without clopidogrel administration. n=5 mice for control, and PLX5622-treated group, and n=6 mice for PLX5622+clopridogrel group. Data are represented as mean ± SEM (bar graphs). Unpaired Student’s t test for (b, d-e, m, o-p), one-way ANOVA with Bonferroni’s post hoc test for (h-k, r-s), *p < 0.05, **p < 0.01, and ***p < 0.001.

Since acute PLX5622 treatment (for 24-48 h) also induces significant upregulation of *Cyp2b10* **(Fig 3i, m, Extended Data Fig. 5g-h)**, we further tested whether CYP2B10 induction accelerates anesthetic emergence. We intraperitoneally injected clopidogrel, a potent CYP2B10 inhibitor impermeable to the blood brain barrier ^53^, into the PLX5622-treated mice to block CYP2B10 activity and measured their responses to pentobarbital anesthesia **(Fig. 3q)**. Blocking CYP2B10 activity largely restored the time of RORR and abolished early arousal from pentobarbital anesthesia after PLX5622 treatment **(Fig. 3r-s)**. Together, these data demonstrate that PLX5622 treatment accelerates anesthesia primarily through induced CYP2B10 activity.

### PLX5622 treatment extensively induces hepatic metabolic enzymes

To further examine the effects of PLX5622 treatment in the body, we profiled the transcriptomes of brain and liver tissues from WT and PLX5622-treated mice **(Fig. 4a, Extended Data Fig. 6a)**. Consistent with microglial elimination, PLX5622 treatment substantially downregulated the expression of microglia-specific genes (e.g. *cx3cr1, csf1r and fcrls*) and the innate immune responses-related pathways (GO: 0002443, 0002253, 0002757) in the brain (**Fig. 4b-c, Extended Data Fig. 6b**). By contrast, PLX5622 substantially induced the expression of drug-metabolizing enzymes in the liver, along with reduced Kupffer cell-associated immune responses (**Fig. 4d-e**, **Extended Data Fig. 6c,**). PLX5622-induced hepatic enzymes are engaged in all three phases of xenobiotic metabolism (Phase I: oxidation; Phase II: conjugation; Phase III: transport) **(Fig. 4f-g)**. Among them, *cyp2b10* and multiple *cyp2c* family genes (*2c29, 2c37,* and *2c54)* are the top induced ones **(Fig. 4d)**. These data demonstrate that PLX5622 treatment induces prominent hepatic changes.

**Fig 4.**
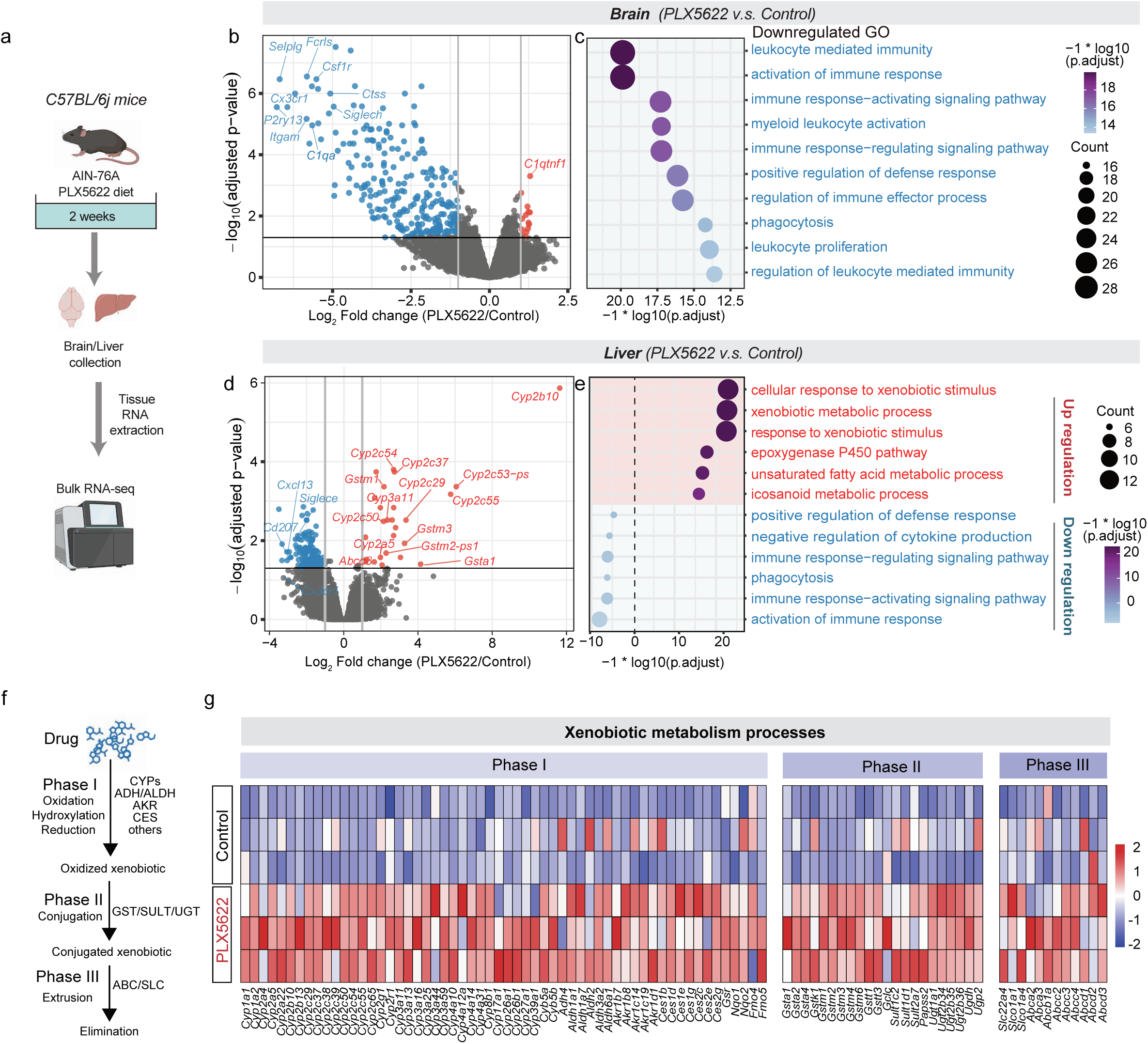
PLX5622 induces robust changes in xenobiotic and endobiotic metabolic pathways in the liver. **(a)** An experimental schedule for RNA-seq. **(b, d)** Volcano plots of transcripts in the brain **(b)** and liver **(d)** from control and PLX5622-treated mice. Red dots indicate upregulated genes, whereas blue dots indicate downregulated genes. **(c, e)** GO analysis of downregulated genes in the brain **(c)** and upregulated genes in the liver **(e)** following PLX5622-treatment. **(f)** Schematic of xenobiotic metabolism pathway highlighting the three phase and related enzyme family. **(g)** Heatmap of xenobiotic metabolism-pathway genes in the liver from the control and PLX5622-treated mice.

### PLX5622 treatment broadly affects xenobiotic and endobiotic metabolism

Hepatic enzymes have profound impacts on xenobiotic and endobiotic metabolism ^54, 55^. To assess whether PLX5622 treatment broadly affects xenobiotic metabolism, we tested several substances ranging from the anti-viral drug-efavirenz **(Fig 5a)**, a known substrate of CYP2b10, to addiction substances such as ethanol and nicotine. We found that PLX5622 substantially enhanced the biotransformation of efavirenz ^56, 57^, manifested by pronounced plasma reduction of efavirenz and concurrent elevation of 8-hydroxy efavirenz, compared to controls **(Fig. 5b-c**). In addition, PLX5622 treatment also promoted ethanol and nicotine metabolism **(Fig. 5h, l-n),** likely due to the induction of CYP family members and alcohol, nicotine metabolism-related genes such as *Aldh1a1*, *Aldh1a7* and *Gstm1*, or *Cyp2a5*, *Cyp2b10*, *Aox1* and *Ugt1a5* **(Fig. 5f, j-k)**. By contrast, their metabolism remains largely unchanged in either DT-treated **(Fig. 5d-e**) or *CSF1R-cKO* mice **(Fig. 5i, o-p**), which are absent of microglia **(Fig. 5 d-e, i, o-p**). Furthermore, levels of plasma cholesterols and triglyceride were also reduced in mice treated with PLX5622 for one-weeks (**Fig. 5q-r**), but not in *CSF1R-cKO* mice (**Fig. 5s-t**), suggesting enhanced endobiotic fatty acid metabolism after PLX5622 treatment. Collectively, these data suggest that PLX5622 treatment extensively modifies the metabolism of a broad spectrum of xenobiotic and endogenous substances, irrelevant of microglia.

**Fig 5.**
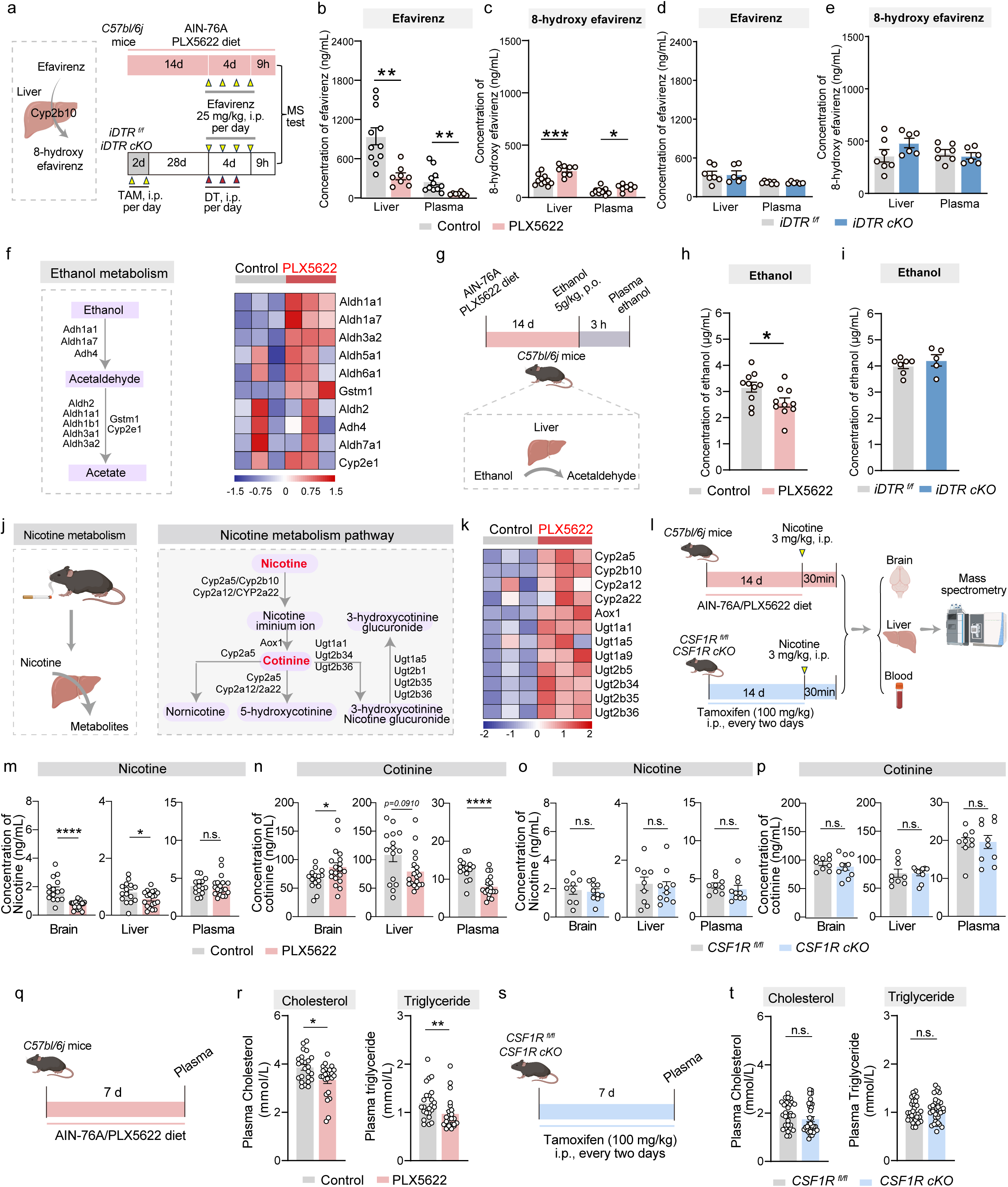
PLX5622 broadly affects xenobiotic and endobiotic metabolism. **(a)** An experimental scheme. **(b-c)** The concentration of efavirenz **(b)**, and 8-hydroxy efavirenz **(c)** in the plasma of control and PLX5622-treated mice. n=11 mice for controls, and n=8mice for PLX5622 group. **(d-e)** The concentration of efavirenz **(d)**, and 8-hydroxy efavirenz **(e)** in the plasma of *iDTR^f/f^*, *iDTR cKO* mice. n=7 mice for *iDTR^f/f^*, and n=6 mice for *iDTR cKO* group. **(f)** Heatmap of ethanol metabolism-pathway genes in the liver from the control and PLX5622-treated mice. **(g)** An experimental scheme. **(h-i)** The concentration of ethanol in the plasma of PLX5622-treated mice **(h)**, and DT-treated *iDTR-cKO* mice **(i)**. n=10 mice for controls and PLX5622 group, n=7 mice for *iDTR^f/f^*, and n=5 mice for *iDTR cKO* group. **(j-k)** Heatmap of nicotine metabolism-pathway genes in the liver from the control and PLX5622-treated mice. **(i)** An experimental scheme. **(m-p)** The concentration of nicotine and cotinine in the brain, liver, and plasma of PLX5622-treated mice **(m-n)** and *CSF1R cKO* mice **(o-p)**, n=15 mice for control mice, and n=19 mice for PLX5622-treated mice, n=9 mice for *CSF1R^f/f^*, and n=10 mice for *CSF1R cKO* group. **(q, s)** An experimental scheme. **(r, t)** The concentration of cholesterol and triglyceride in the plasma of PLX5622-treated mice **(r)** and *CSF1R cKO* mice **(t)**, n=24 mice for control mice, and n=23 mice for PLX5622-treated mice, n=30 mice for *CSF1R^f/f^*, and n=33 mice for *CSF1R cKO* group. Data are represented as mean ± SEM (bar graphs). Unpaired Student’s *t* test for **(b-e, h-i, m-p, r, t)**. \**p* < 0.05, \*\**p* < 0.01, and \*\*\**p* < 0.001.

### PLX5622 treatment alleviates nicotine withdrawal-induced anxiety phenotype independent of microglia

PLX5622 has also been used to investigate the role of microglia in the contexts of addiction ^19, 22, 23, 25^. Reduced addictive signs and addiction abstinence have been observed in multiple models, including nicotine addiction ^19, 21, 25^, and microglial contribution had been proposed. However, as PLX5622 treatment also significantly promoted nicotine metabolism, it is likely that reduced phenotypes may be due to enhanced nicotine metabolism, rather than microglial depletion.

To assess whether microglia regulate nicotine withdrawal behaviors, we administered nicotine to control, PLX5622-treated, or *Cx3Cr1^CreER/+^::CSF1R^fl/fl^*mice (after tamoxifen induction), and analyzed anxiety-related behaviors upon nicotine withdrawal **(Fig. 6a, e)**. Consistent with previous reports ^19, 21, 25^, PLX5622-treatment significantly reduced anxiety upon nicotine withdrawal **(Fig. 6b-d)**. By contrast, genetic depletion of microglia in *Cx3Cr1^CreER/+^::CSF1R^fl/fl^*mice showed no differences in anxiogenic behaviors from controls **(Fig. 6f-h)**. Together, these data suggest that PLX5622 treatment-induced anxiety alleviation upon nicotine withdrawal results from enhanced nicotine metabolism, rather than microglial regulation.

**Figure 6.**
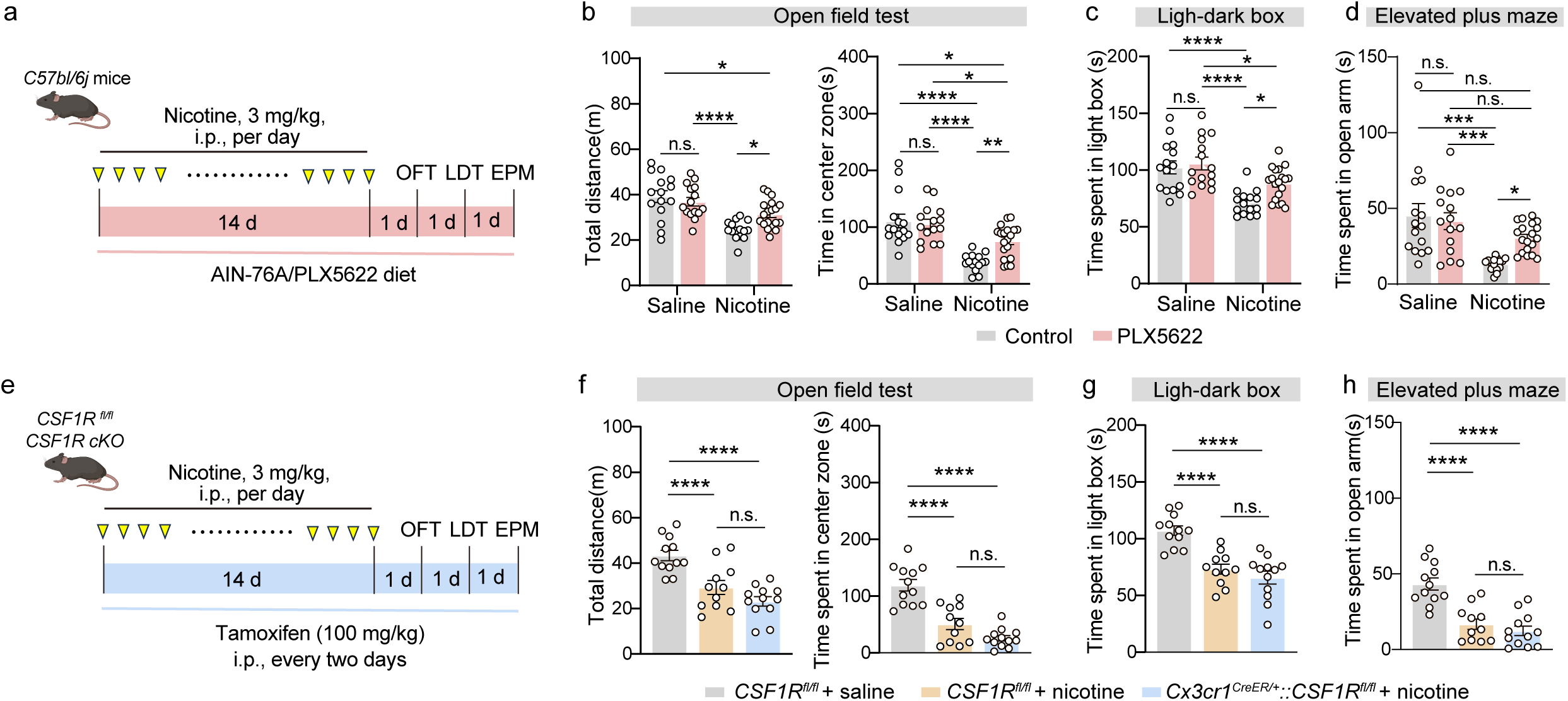
PLX5622 treatment alleviates nicotine withdrawal-induced anxiety phenotype independent of microglia. **(a, e)** An experimental scheme. **(b-d, f-g)** Open field test, light-dark box test, and elevated plus maze test after saline or nicotine withdrawal between control and PLX5622-treated mice **(b-d)** or *CSF1R^f/f^*, *CSF1R cKO* group **(f-g)**. n=15 mice for control-saline group, n=15 mice for PLX5622-saline group, n=15 mice for control-nicotine group, and n=19 mice for PLX5622-nicotine group; n=12 mice for *CSF1R^f/f^*-saline group, n=11 mice for *CSF1R^f/f^*-nicotine group, n=12 mice for *CSF1R cKO* -nicotine group. Data are represented as mean ± SEM (bar graphs). One-way ANOVA with Bonferroni’s post hoc test for (**f-h**), two-way ANOVA with Bonferroni’s post hoc test for (**b-d**), \**p* < 0.05, \*\**p* < 0.01, and \*\*\**p* < 0.001.

## Discussion

Diverse functions of microglia have been revealed using CSF1R inhibitors. While studies have optimistically linked different phenotypes to microglial depletion, here, we uncover that PLX5622, a widely used CSF1R inhibitor, significantly induces metabolic enzymes in the liver and extensively modifies xenobiotic and endobiotic metabolism. Such systemic metabolic reprograming generates confounders in using PLX5622 to reveal microglial function in both anesthesia and addiction models. Our work provides foundation for rigorous evaluation of experimental results using PLX5622 to understand microglial biology and presents new insights for strategic design of next-generation inhibitors.

Microglia intimately interact with neuronal elements to fine-tune neuronal activity ^3, 35, 36^, and microglia-mediated neuronal activity regulation has been shown to be involved in the pathogenesis of epilepsy, and stroke ^2, 3, 58^. Recent observations that PLX5622 treatment shortened injectable anesthetic duration have prompted a proposal that microglia maintain and facilitate anesthesia by balancing the neuronal network between anesthesia-promoting and anesthesia-inhibiting neurons. In further characterizing microglial contribution in anesthesia, we were unable to observe significant changes in neuronal activity or extracellular adenosine in the anesthetized brain after PLX5622 treatment. However, we noticed that PLX5622 treatment selectively affects injectable anesthesia, with no apparent effects on inhalation anesthesia and sleep. The universal and dramatic effects of PLX5622 treatment on all tested injectable anesthesia are contrasted with no measurable influence on inhaled anesthesia or sleep. Moreover, PLX5622 treatment had much stronger influences than DT treatment in promoting anesthetic emergence ^7^. Although this discrepancy can be partly explained by the differences in drug potency and efficacy of microglial depletion, it is puzzling that PLX5622 even outperformed direct neuronal manipulation in accelerating recovery ^40, 59, 60^. All of these data point out that, alternative mechanisms, other than microglia-medicated neuronal control, may be involved in these interesting phenotypes.

Our findings that PLX5622 treatment prominently enhanced the metabolism of injectable anesthetics clarify these conflicting observations. These data are further supported by the discovery that PLX5622 prominently induced the expression of a number of metabolic enzymes, which help promote the biotransformation and elimination of anesthetics. These data suggest that PLX5622 treatment-resulted shortened anesthesia are mainly driven by enhanced drug metabolism, which reduced the drug efficacy. The fact that DT treatment-induced genetic ablation of microglia or microglial P2RY12 deficiency also shortened anesthetic duration suggest that microglia indeed tune anesthesia. Such tuning may be subtle and intricate that requires further efforts to find out the underlying mechanisms.

PLX5622 is touted as being a highly specific CSF1R inhibitor ^9^. However, we demonstrate that feeding mice with the standard dose of PLX5622 (1,200 mg/kg) in chow for one week could potently induce the expression of CYP2B10, by more than 7000 times in the liver. Other than CYP2B10, PLX5622 also pronouncedly upregulated a large repertoire of hepatic enzymes, which are involved in the disposition of a broad range of xenobiotic drugs, as well as endogenous metabolites. Indeed, PLX5622 treatment substantially modified the metabolisms of different drugs, including nicotine, cocaine and morphine (**Extended Data Fig. 6D-E**), which also produces confounding effects in nicotine-withdrawal phenotypes. Moreover, the reduction of plasma cholesterol and triglyceride in PLX5622-treated mice suggests that PLX5622 may modify endobiotic fatty acid metabolism. Consistent with this, PLX5622 has been shown to alters the cholesterol metabolism in brain endothelial cells ^29^. The profound impacts of PLX5622 on xenobiotic and endobiotic metabolism raise awareness regarding the application of PLX5622 in studies to avoid potential drug-drug interactions, particularly in the context of drug-induced models or metabolic diseases.

In summary, in uncovering the mechanisms underlying microglial contribution in anesthesia, we serendipitously found that PLX5622 induced pronounced hepatic effects that confounds understanding microglial functions during anesthesia. These previously unrecognized findings raise important concern and awareness when using PLX5622 for scientific investigation and underscore the importance of combining different approaches for accurate evaluation of microglial functions. Our study also provides valuable insight into strategic designing of selective CSF1R inhibitors for translational therapy-related application in cancer and neurodegenerative diseases.

## Methods

### Animals

C57BL/6j mice (7-16 weeks of age, 20-30 g) were purchased from Charles River (China). *Cx3cr1^CreERT2^* mice were purchased from the Jackson Laboratory (Jax #: 021160). *R26^iDTR/iDTR^*mice (Jax #: 007900) were obtained from Dr. Hailan Hu (Zhejiang University, China), *CSF1R^fl/fl^* mice (Jax #: 021212) were obtained from Dr. Jiyun Peng (Hangzhou Normal University, China). All animals were maintained in the Animal Core Facility of Zhejiang University under a 12h light/dark cycle with free access to food and water. All animal experimental protocols were approved by the Animal Care and Use Committee of the animal facility at Zhejiang University.

## METHOD DETAILS

### Drugs and injection

For induction of the Cre recombinase, tamoxifen (150 mg/kg, Sigma) were given to the adult *Cx3cr1^CreERT2/+^::R26^iDTR/iDTR^* mice by intraperitoneal injection at 48-hour intervals, and tamoxifen (100 mg/kg) were given to the *Cx3cr1^CreERT2/+^:: CSF1R^fl/fl^* mice by intraperitoneal injection at 48-hour intervals for a total of 14 days. Three doses of diphtheria toxin (DT, Cat# 150, List Biological Laboratories Inc, 0.03 mg/kg per dose) dissolved in normal saline were intraperitoneally administered every 24 hours. To measure the metabolic activity of CYP2B10, efavirenz (25 mg/kg, TargetMol, USA), a known substrate of CYP2B10, was administered to the animals by oral gavage for four consecutive days and sacrificed by decapitation 9 h after the last administration according to previous description. To measure the metabolic rate of sodium pentobarbital, mice were intraperitoneally injected with sodium pentobarbital (100 mg/kg) and anesthetized for 10 min, 1 h, and 2 h as indicated.

### PLX5622 treatment

To effectively ablate microglia, adult mice were continuously fed with PLX5622-containing chow (1, 200 mg/kg, Plexxikon Inc., Berkeley) for two weeks and control mice were fed with normal AIN-76A diet ^9^.

### Immunofluorescence analysis

Brains or livers were dissected and post-fixed in 4% paraformaldehyde (PFA, Sigma-Aldrich), dehydrated in 30% sucrose solution, embedded in the optimal cutting temperature (OCT, SAKURA Tissue-Tek) solution and sliced at a thickness of 30 μm using a cryostat microtome (Lecia CM1860 UV, Japan). Brain and liver slices were antigen-retrieved in the citrate buffer (10 mM sodium citrate, 0.05% Tween-20, pH 6.0), permeabilized in 0.5 % Triton X-100 for 10 minutes, and blocked in 5% (wt/vol) bovine serum albumin for 1 h at room temperature, followed by incubation with primary antibodies as indicated (rabbit anti-Iba1, 1:1000; Rat anti-cFos, 1:5000) overnight at 4 °C. Slices were washed three times in Tris-buffered saline containing 0.5% Tween-20 (TBS-T) and incubated with secondary antibodies, followed by TBS-T washes. Nuclei were counterstained with DAPI (Beyotime, China) and slides were mounted using anti-fade reagents (Millipore, USA). The immunofluorescence images were captured by an FV-1200 confocal microscope (Olympus, Japan). For cell counting, images were obtained from at least three animals, and the numbers of cells from at least three slices of each animal were counted by CellSens Dimension (Olympus, Japan).

### c-Fos screening

Mice were fed with control or PLX5622 diet for 2 weeks and gently handled for 3-4 days before the experiment. Brains were dissected 90 min after intraperitoneal injection of 0.2 mg/kg dexmedetomidine and 100 mg/kg pentobarbital. Immunostaining for c-Fos were conducted as mentioned before.

### Analyses of general anesthesia (GA)

For injectable anesthesia, mice were placed into an acrylic glass chamber for 15 min for environmental adaptation, followed by intraperitoneal injection of sodium pentobarbital (50 or 100 mg/kg).

For inhalation anesthesia, mice were placed into an acrylic glass chamber connected to an isoflurane (RWD, China) or sevoflurane (RWD, China) vaporizer 10 min before isoflurane or sevoflurane administration, then the drug was given to the chamber with a 1.0 L/min flow of 100% oxygen. After a 30 min exposure to isoflurane (1%, 1.5%, 2%) or sevoflurane (3%), the chamber lid was open and isoflurane or sevoflurane was replaced by room air.

The responses of mice during the whole period were video-recorded and the time of anesthetic induction (loss of righting reflex, LORR) and recovery (recovery of righting reflex, RORR) were calculated by an investigator blinded to the experimental conditions. LORR was defined as the state failed to right themselves for at least 15 s when lying on their backs, and checked every 10 s. Turning over by themselves with four paws standing on the floor were considered to exhibit RORR. The interval between LORR to RORR was defined as anesthetic duration.

### Stereotaxic Surgery for placement of electroencephalogram (EEG) and electromyogram (EMG)

All mice were anesthetized with sodium pentobarbital (100 mg/kg) (i.p. injection) and placed in a stereotaxic frame (David Kopf Instruments). A customized EEG and EMG unit was implanted on the skull of the mice, with two integrated EEG electrodes and two EMG electrodes inserted into the neck muscles at the rear of the skull, as described previously. The four stainless-steel anchorage screws of the unit were fixed to the skull with dental cement, and tissue glue (3 M Vetbond, St. Paul, MN, USA) was applied to help the wounds heal and the electrodes fixed. After the surgery, all animals were allowed to recover in individual chambers for 1 week, and were handled daily for 10 min for habituation. To allow free movement in the EEG/EMG recording chamber without tangling, a slip-ring device (CSF-22, Biotex, Japan) was used for the cable. After that, each animal was transferred to an EEG/EMG recording chamber and connected to an EEG/EMG head-stage. The animals were habituated for 2 days before recording.

### EEG/EMG recording and analysis

EEG signals were recorded from electrodes on the frontal cortices (AP, 2 mm; ML, 1 mm). The EEG and EMG signals were amplified and filtered (EEG, 0.5 Hz–50 Hz; EMG, 10–500 Hz) by an AC filter, and recorded at a sampling rate of 200 Hz (PowerLab ML795, ADInstruments, Dunedin, Australia) using the LabChart software (ADInstruments, Dunedin, Australia).

The EEG power spectral density was analyzed using NeuroExplorer (Nex Technology, Littleton, MA) and MATLAB (MathWorks, Cambridge, UK). EEG power spectrum was calculated using fast Fourier transform for the frequency range of 0-30 Hz. Total power percentages of the frequency bands of delta (0.5-4 Hz), theta (4-10 Hz), alpha (10-15 Hz), beta (15-25 Hz) were calculated to evaluate anesthetic states. Time to the loss of consciousness (LOC) and recovery of consciousness (ROC) were determined by EEG and EMG signal. LOC is determined by the onset of strong delta power in EEG and minimal muscle tone in EMG recording; ROC is determined by the reduction of delta power in EEG and continuously-elevated muscle activity for more than 1 min.

### Sleep-waking state classification

Sleep states were auto-scored using sleep analysis software (SleepSign, Kissei Comtec, Japan) on the basis of the EEG and EMG waveforms in 4 s epochs. Different sleep states were defined as follows: wakefulness, desynchronized EEG with high EMG activity; NREM, synchronized EEG with high-amplitude of low frequencies (0.5-4 Hz, delta power) and low EMG activity; REM, desynchronized EEG with high power at theta frequencies (4-10 Hz) and low EMG activity. All classifications of states assigned by SleepSign were examined visually and corrected manually.

### Stereotaxic injection, fiber implantation and photometry recording

Mice aged 8-10 weeks were deeply anesthetized by intraperitoneal injection of pentobarbital (100 mg/kg). The mice were fixed in a stereotaxic apparatus with the skull aligned. After sterilization, the scalp was opened and a small hole was drilled in the skull according to the targeted coordination. For fiber photometry recording, 200 nL AAV2/9-hSyn-GRAB_Ado_ 1.0 virus (provided by Yulong Li lab at Peking University) was injected into the mPFC (A/P +1.97 mm, M/L +0.4 mm, D/V −2.00 mm) of mice at 40 nL/min via a Hamilton syringe. Then, an optical fiber (200 μm in diameter) was implanted 50 μm above the injection site and was secured with dental cement. All experiments were carried out at least two weeks after surgery for virus expression.

Two weeks after the surgery, an optic fiber was attached to each implanted ferrule via a ceramic sleeve for fluorescence recording. The GRAB sensor was excited by 470 nm excitation light, and the emission fluorescence was recorded using a two-color fiber photometry system (Inper LLC). The 410 nm excitation light was delivered in the same patch cord and used to correct bleaching and noise fluctuations due to animal movements. Data were collected at an exposure time of 20 ms and at a sampling rate of 25 Hz.

Data were analyzed using Inper Analysis software (Inper plot, Inper LLC). For the average plots, the onset of isoflurane-inhalation or ketamine-injection was aligned to the time zero and the changes of fluorescence (ΔF/F) were calculated as (F-F0)/F0, where F0 was the mean fluorescence signal from −300 s to 0 s. The heatmaps illustrating the fluorescence changes (ΔF/F) for each mouse individually was generated from MATLAB using a customized script.

### RNA extraction and quantitative RT-qPCR

Total RNA from brain and liver tissues were extracted using TRIzol (Takara Bio, USA). RNA was reverse-transcribed using PrimeScript RT Master Kit (Takara Bio, USA). Real-time PCR was performed using the TB Green Premix Ex Taq™ (Tli RNaseH Plus) on a Bio-Rad CFX96 Real-Time PCR Detection System (Bio-Rad, USA). Primer sequences are provided in the **Supplementary Table 1**. The relative expression was measured using the 2^−ΔΔCt^ method. ΔCt = Ct _(Target genes)_ – Ct_β-actin_. ΔΔCt = ΔCt_(Target genes)_ – ΔCt _(average ΔCt of control)_. Relative fold changes were determined by 2^−ΔΔCt^ and normalized to the expression levels of β-actin.

### Western blotting

Liver tissue was lysed in RIPA lysis buffer (Beyotime) by sonication and protein extracts were obtained from the supernatants after centrifugation. The extracts were subjected to sodium dodecyl sulfate-polyacrylamide gel electrophoresis and electro-transfer. The membranes were blocked with 5% BSA, followed by incubation with primary antibodies and species-appropriate HRP-conjugated secondary antibodies. Membranes were subsequently washed in PBS-T three times and immunodetected with enhanced chemiluminescence reagents (PerkinElmer).

### Antibodies

All primary and secondary antibodies with source, dilutions and validations are listed in **Supplementary Table 2.**

### RNA sequencing

Mouse brain and liver tissues were harvested and total RNAs were extracted using TRIzol (Takara Bio, USA). The purity, concentration, and integrity of RNA were evaluated using the RNA Nano 6000 Assay Kit from the Agilent Bioanalyzer 2100 system. Quality control, cDNA library generation, and subsequent RNA sequencing services were performed by Novogene corporation (Beijing, China). Briefly, oligo (dT) beads were utilized to enrich mRNA, which was then fragmented. The cDNA libraries were generated using NEBNext Ultra RNA Library Prep Kit for Illumina. The purified and processed cDNA libraries were assessed for insert size using the Bioanalyzer 2100, for concentration using the Qubit assay, and for quality using qPCR. The cDNA library samples were sequenced on the Novaseq6000 with a paired-end 150 configuration. Raw reads were filtered and aligned to the mouse genome (Mus musculus GRCm38, NCBI). Based on these read counts, normalization and assessment of differential gene expression were performed using DESeq2 on R (4.2.0). Genes with fragments per kilobase million lower than 1 (FPKM < 1) in all samples were excluded from the subsequent analyses. Differential expressed genes (DEGs) were analyzed using the R package bases on the raw counts, with a fold change greater than 2 and adjusted P value less than 0.05 in comparison to the control group. Volcano plots and heatmaps were visualized by R package (ggplot2 and gplot), and GO enrichment was visualized by Bioconductor package (“cluster profiler” package).

### Sample preparation and mass spectrometry analyses

Blood samples were collected from mice treated with pentobarbital or efavirenz in heparin sodium-coated tubes, followed by centrifugation at 5, 000 rpm for 10 min at 4° C. The resulting plasma supernatants were separated and stored at −80°C until further analysis. After blood collection, mice were perfused with ice-cold normal saline. Brains and livers were dissected, snap-frozen in liquid nitrogen, and stored at −80°C until analysis

For tissue sample preparation, 100 mg of brain or liver were weighted, and 0.15 ml ddH_2_O was added for homogenization using a tissue grinder at 60 Hz for 3 min (JXFSTPRP-24L, Jingxin, Shanghai, China). Next, 450 μl acetonitrile was added to the homogenized samples, and thoroughly vortex mixed, and placed at 4 °C overnight. The mixture was centrifuged at 12,000 rpm for 30 min at 4 °C, and the supernatants were filtered through a 0.22 μm membrane. A 70 μl aliquot of the filtered supernatants was then injected into a Waters Alliance 2795 HPLC system (Waters Corp., Milford, MA, USA) coupled to a Waters Quattro Premier mass spectrometer (Waters Corp., Milford, MA, USA).

The HPLC autosampler was maintained at 10°C, and chromatographic separation was performed using a Waters Acquity UPLC BEH C18 column (2.1 mm × 50 mm inner diameter, 1.7 μm particle size; Waters) or ZORBAX SB-AQ Rapid Resolution HT column (4.6mm × 50 mm inner diameter, 1.8 μm particle size, 600 bar; Agilent) at 30° C.The mobile phase consisted of pure methanol (component A) and 0.1% ammonium acetate buffer (component B) at a flow rate of 0.2 ml/min with an injection volume of 5 μl. A linear gradient of tested chemicals, including pentobarbital, efavirenz, and 8-hydroxy efavirenz, and nicotine were employed. For MS/MS detection, electrospray ionization in negative (ESI−) or positive (ESI+) mode was used in multiple reaction monitoring (MRM) mode, with a dwelling time of 0.1 s per transition. The ESI source was set with the following conditions: capillary voltage of 3.0 kV, source temperature of 120 °C, desolvation temperature of 350 °C, cone gas flow of 50 L/h, and desolvation gas (nitrogen) flow of 800 L/h. Specific m/z transitions, collision energies, and retention times for all compounds are provided in the. Data acquisition and peak integration were performed using MassLynx 4.0 software with QuanLynx v4.1 (Waters®). A standard calibration curve was generated for each analyte using different concentrations, and the concentrations of the chemicals in the injected samples were quantified based on the area ratios.

### High-resolution mass spectrometry analyses

Liver samples were collected from control and PLX5622-treated mice 1 hour after pentobarbital injection. Tissues were homogenized, followed by protein precipitation and centrifugation to obtain the supernatants. After vacuum drying the supernatants, the residue was resuspended in 800 μL of 20% acetonitrile. The sample was agitated at 20 Hz for 10 minutes and incubated on ice for 30 minutes. Following incubation, the mixture was centrifuged three times at 15,000 g for 15 minutes at 4 ° C. Chromatographic separation was performed using a Hypersil GOLDTM C18 column (2.1 mm × 100 mm inner diameter, 1.9 μm particle size) maintained at 40 °C. The mobile phase consisted of 0.1% formic acid in water (Solvent B) and 100% acetonitrile (Solvent A) with a flow rate set to 0.3 ml/min. The injection volume was 1 μl. A linear gradient elution profile was applied as follows: 0–1 min, 10% A; 1–5 min, linear increase from 10% to 90% A; 5–7 min, 90% A; 7–7.01 min, linear decrease from 90% to 10% A; and 7.01–10 min, 10% A for re-equilibration. Mass spectrometric detection was carried out using an Orbitrap Exploris 240 MS (Thermo Scientific) equipped with a heated electrospray ionization (ESI) source. The MS parameters for negative ion mode included a spray voltage of −2.5 kV, and ion source, capillary, and auxiliary gas temperatures of 320 °C, 300 °C, and 300 °C, respectively. The gas flow rates were set to 8, 1, and 40 arbitrary units (AU) for auxiliary, sweep, and sheath gases, respectively. For qualitative analysis, a full MS/data-dependent MS2 (ddMS2) acquisition strategy was employed. The automatic gain control (AGC) target value was set to 1.0×10^6 with a maximum injection time of 100 ms. The full MS scan range was set to m/z 70– 1050 with a resolution of 60, 000. The S-lens RF level was adjusted to 35 AU. For ddMS2 acquisition, product ion spectra were acquired at a resolution of 15000 with a mass isolation window of 1.5 m/z. The RF lens was adjusted to 70 AU. Thermo Xcalibur™ 4.4 was used for data acquisition and interpretation.

To confirm the identification of pentobarbital and 3-hydroxy pentobarbital, LC-MS/MS with a triple quadrupole mass spectrometer was employed (Triple Quad 5500+ System; AB SCIEX). Chromatographic separation was performed using an ACQUITY UPLC® HSS T3 column (2.1 mm × 150 mm inner diameter, 1.8 μm particle size) maintained at 40 °C. The mobile phase consisted of 0.1% formic acid in water (Solvent B) and 100% acetonitrile (Solvent A) with a flow rate set to 0.3 ml/min. The injection volume was 1 μl. The raw data were analyzed using OS v2.0 software.

### Determination of the ethanol Using Gas Chromatography (GC)

Blood was collected from control and PLX5622-treated mice 3 hours after ethanol administration (oral gavage in a volume of 0.02 mL/g to a final concentration of 5 g/kg), followed by centrifugation at 5000 rpm for 10 min at 4°C to obtain plasma. The concentrations of ethanol were quantified using a gas chromatography (GC) method equipped with a flame ionization detector (FID). The analysis was performed using a SHIMADZU Nexis GC-2030 system coupled with a SH-Rtx™-624 capillary column (20 m × 0.18 mm i.d., 1.0 µm film thickness). A 50 µL plasma sample was transferred into a 1.5 mL Eppendorf tube, followed by the addition of 100 µL of dichloromethane (DCM). The mixture was vortexed briefly and then stored at −80°C for 3 hours to facilitate phase separation. After incubation, the sample was centrifuged at 12,000 rpm at 4°C for 20 minutes. The supernatant was carefully collected and used as the test solution for subsequent GC analysis.

For chromatographic separation, a temperature program was employed as follows: the initial oven temperature was set at 50°C and held for 2 minutes, followed by a ramp of 10°C/min to 100°C, where it was maintained for 1 minute. A split ratio of 50:1 was applied to control sample loading, with nitrogen gas serving as the carrier gas. The volumetric flow rate through the column was maintained at 1.94 mL/min.

Ethanol was identified based on its characteristic retention times by comparing it to authentic standards of anhydrous ethanol (both prepared at 1% concentration in isopropanol). Quantification was carried out using the G1701BA Chem Station B.01.00 software (Agilent Technologies, Alpharetta, GA, USA), which calculated the relative concentrations of ethanol in plasma samples by determining the ratio of the standard peak area to the peak area of the analyte of interest.

### Biochemical analyses of plasma

Blood samples from WT, PLX5622-treated mice, *CSF1R^fl/fl^* and *Cx3cr1^CreER/+^::CSF1R^fl/fl^* mice were collected and transferred into heparin sodium-coated tubes, followed by centrifugation at 5000 rpm for 10 min at 4°C. The resulting plasma supernatants were was carefully separated and stored at −80°C prior to use. Plasma levels of triglyceride (TG) and total Cholesterol (TC) were detected using the TG Enzymatic Determination Kit (A111-2-1, NanJing JianCheng Bioengineering Institute, Nanjing, China) and the TC Enzymatic Determination Kit (A110-2-1, NanJing JianCheng Bioengineering Institute, Nanjing, China).

### Nicotine-induced addiction model

(−)-Nicotine tartrate (Cayman Chemicals, USA) was dissolved in 0.9% sterile sodium Chloride solution (pH 7.0) and administered to mice at a dose of 3 mg/kg. Mice received intraperitoneal (i.p.) injections twice daily at 5-hour intervals for 14 consecutive days. hronic nicotine treatment at this dose results in a plasma concentration of approximately 12.5 nM (reported as nicotine free base molecular weight), which is comparable to the plasma levels observed in human smokers consuming 1–2 cigarettes per day (3.5–18 nM)(REF: Guidelines on nicotine dose selection for in vivo research. Psychopharmacology 190, 269–319 (2007)). Control animals received i.p. injections of 0.9% sodium chloride solution only. Following 14 days of chronic administration with either nicotine or saline, mice in the withdrawal groups underwent spontaneous withdrawal by cessation of the respective treatment.

### Behavioral analysis

#### Open Field test

OFT was conducted before all other behavioral tests. Locomotion and anxiety tests were assessed by modified version of the open field test in a square open field (55 cm length × 50 cm width × 50 cm height). All mice were tested at the 24-hour withdrawal time point. Mice were placed individually into the center of the field and allowed to explore for 10 min. The activity of mice in the field was recorded using a movement tracking system (ANY-MAZE software) connected to a camera mounted above the field. Time spent in the center zone was assayed as a measure of anxiety-like behavior. Locomotor activity was evaluated as the total distance traveled.

#### Light-Dark Box Test

The light-dark box test was used to analyze anxiolytic- or anxiogenic-like behavior in mice. All mice were tested at the 48-hour withdrawal time point. The exterior size of the typical chamber is 57×21×25 cm, which contains a dark compartment (21×15×25 cm), a light compartment (30×21×25 cm), connected by a 6×4×5 cm channel in the middle. For habituation, mice were allowed to move freely within the chamber for 5 min. For the test, individual mouse was placed in the middle of the light compartment, and activities of the animal within the chamber and duration in the light and dark areas were recorded for 5 min using a movement tracking system (ANY-MAZE software).

#### Elevated plus maze test

The maze consisted of four crossing arms (two open and two closed) placed at a height of 50 cm above the ground. All mice were tested at the 72-hour withdrawal time point. Mice were placed in the center of the platform at the beginning of the experiment and allowed to freely explore the maze for 5 min. Locations of mice were tracked automatically by the ANY-maze.

#### Quantification and statistical analysis

Statistical analyses were performed using GraphPad Prism (Version 9.0.0, CA, USA). All quantitative results are presented as the mean ± standard errors of means (SEM) from at least three independent experiments. Statistical differences were analyzed by two-tailed Student’s *t*-test for two groups, or one-way ANOVA with Bonferroni’s post hoc tests for more than two groups. Nonparametric tests were conducted when data are not normally distributed, including Mann-Whitney test. *p*<0.05 was considered significant.

## Lead Contact

Further information and requests for resources and reagents should be directed to the Lead Contact, Zhihua Gao.

## Materials Availability

The study did not generate new unique reagents.

## Data and Code Availability

- The bulk RNA-seq data of liver from C57BL/6j, PLX5622-treated C57BL/6j mice have been deposited at GEO and are publicly available as of the date of publication.
- This paper does not report original code.
- Any additional information required to reanalyse the data reported in this paper is available from the lead contact upon request.

## ACKNOWLEDGMENTS

We are thankful to Wenzhao Zhou at the Core Facilities of Liangzhu Laboratory for mass spectrometry and gas chromatography technical assistance. This work was supported by the National Natural Science Foundation of China (32371003 and 32070974 to Z.G., 32300793 to K.C.); National Science and Technology Innovation 2030-Major Projects (2021ZD0202700 to Z.G.), National High-Level Talent Special Support Programs (10,000 Talents Program) Leading Talents to Z.G, and High-level Innovative Talents in Health Care, Zhejiang Province (519900-52201 to Z.G.), the China Association for Science and Technology (YESS20230262 to K.C.), the Natural Science Foundation Exploration Project of Zhejiang Province (Y24C090012 to K.C.).

## AUTHOR CONTRIBUTIONS

Z.G. supervised the project; Z.G., K.C., W.C. and L.Q conceived the project and designed experiments; K.C., W.C., L.Q., Z.W., Y. Z., Y.Y. and J.X., carried out experiments; K.C., W.C., L.Q. and W.W. analyzed the data; C.Y., J.X., W.Z., and C.M. provide technical assistance, Z.G., K.C., W.C. and L.Q wrote the paper; Z.G., K.C., W.C., L.Q., L.Z., Z. W., H.M., T.H. and C.Y. reviewed and edited the paper; Z.G., and K.C. provided funding acquisition.

## INCLUSION AND DIVERSITY

We support inclusive, diverse, and equitable conduct of research.

---

**Extended Data Fig 1.**
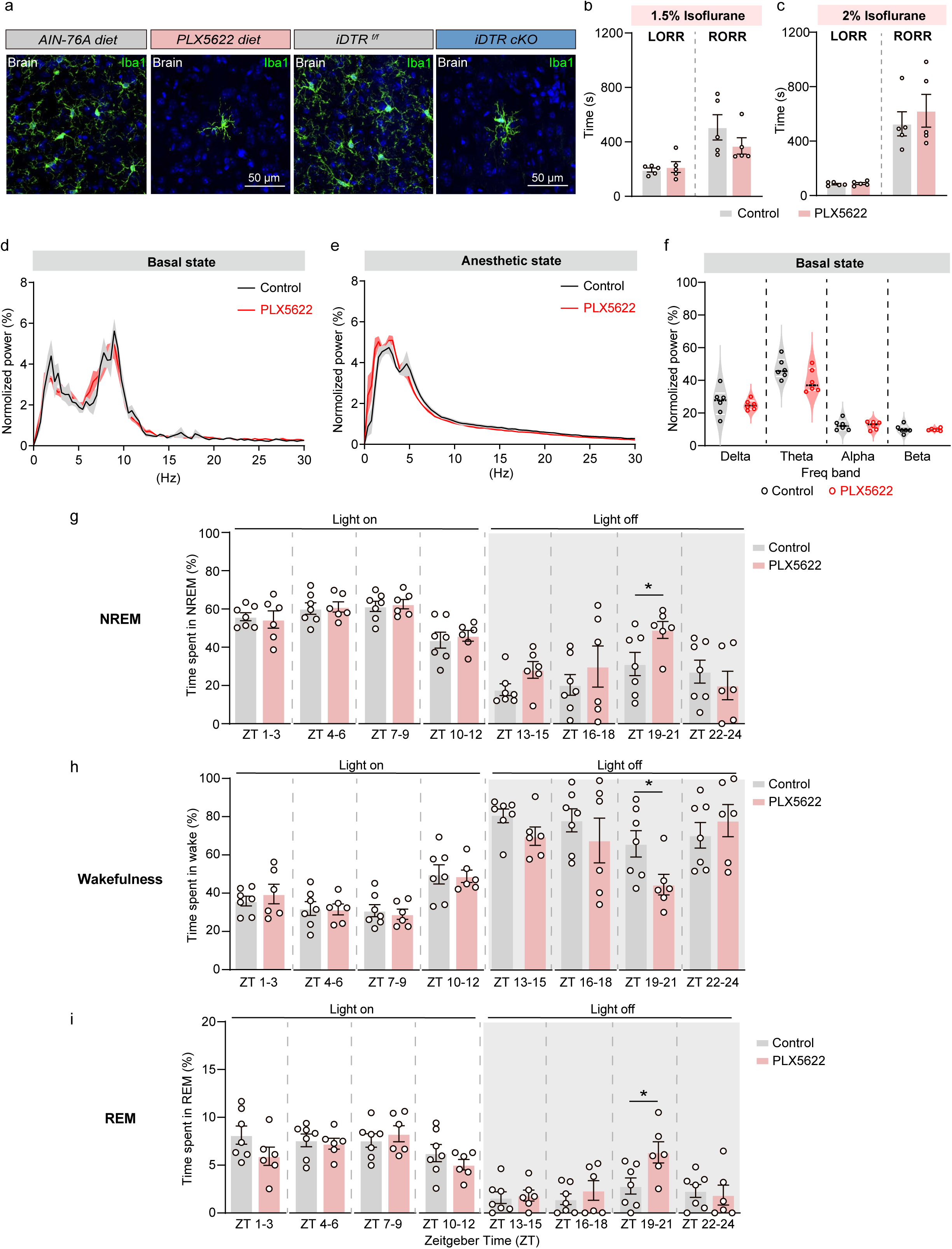
Microglial depletion did not affect isoflurane-induced general anesthesia and slightly altered the sleep patterns. **(a)** Representative images of Iba1 staining from the cerebral cortex in PLX5622 treatment or iDTR cKO mice. Iba1^+^ (green) cells represent microglia. **(b-c)** Time of anesthetic induction (LORR) and recovery (RORR) after 1.5% **(b)**, 2% **(c)** isoflurane exposure between control and microglia-depleted (PLX5622-treated) mice. n=5 mice per group. **(d-e)** Normalized power densities of EEG signals before (d, 15 min of wakefulness) and during anesthesia (e, time periods between LOC and ROC) in control or PLX5622-treated mice. n=6 mice per group. **(f)** Power distribution of EEG frequency bands during wakefulness in control and PLX5622-treated group. **(g-i)** Percentages of NREM (g), wakefulness (h), and REM (i) during sleeps in 3-h bins in control and PLX5622-treated mice. n=7 mice for control and n=6 mice for PLX5622 group. Data are represented as mean ± SEM (bar graphs). Unpaired Student’s *t* test for **(b-c, f-i)**, \**p* < 0.05, \*\**p* < 0.01, and \*\*\**p* < 0.001.

**Extended Data Fig 2.**
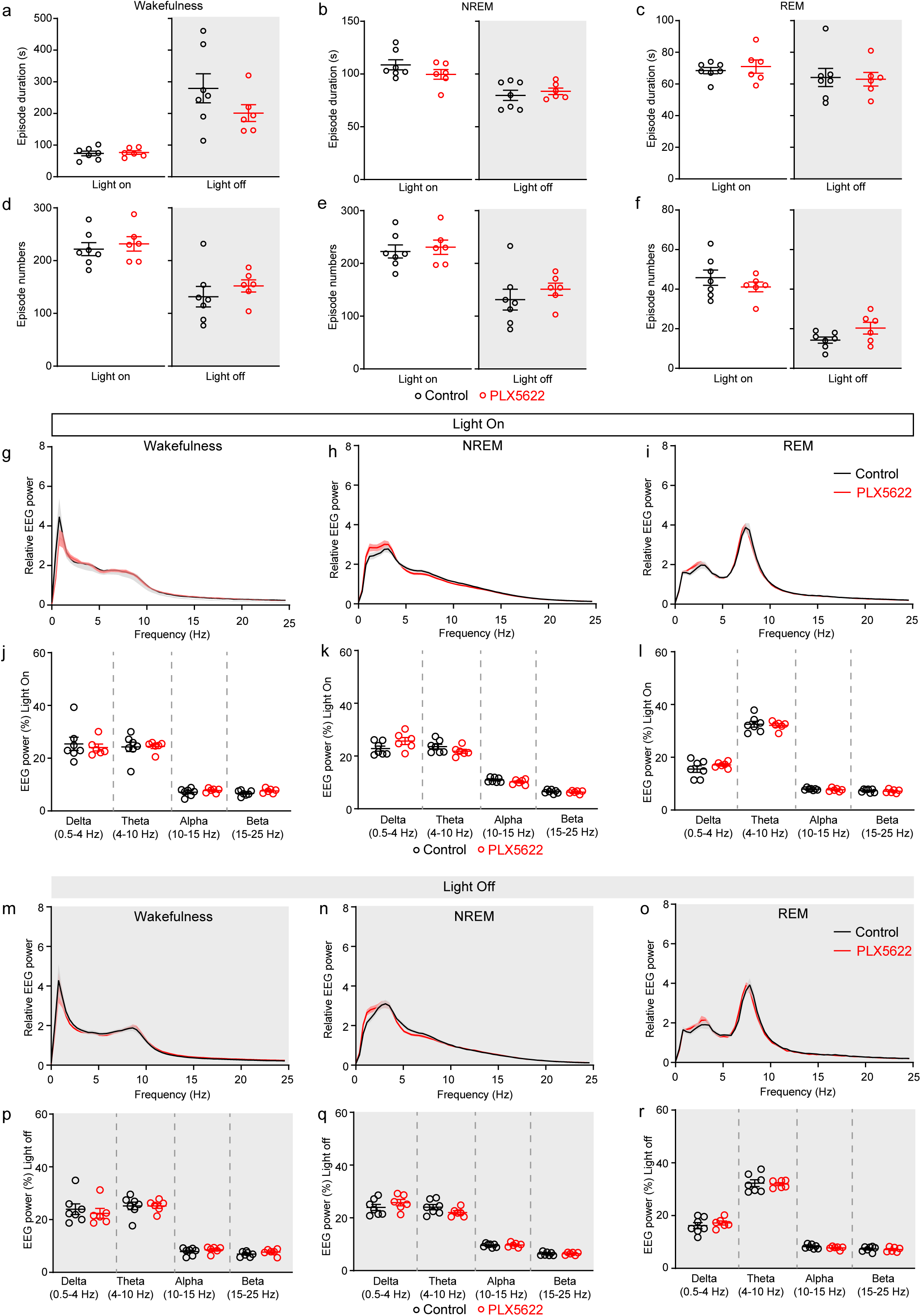
Microglial depletion did not affect the episodes and the EEG power spectrum of wakefulness, NREM and REM sleep. **(a-c)** Episode duration of wakefulness (a), NREM (b) and REM (c) sleep of control and PLX5622 mice across the “light on” and “light off” periods. **(d-f)** Episode numbers of wakefulness (d), NREM (e) and REM (f) sleep of control and PLX5622 mice across the “light on” and “light off” periods. **(g-i)** Normalized EEG power during wakefulness (g), NREM (h) and REM (1) in control and PLX5622 mice across the “light on” periods. **(j-l)** Quantification of EEG power in the delta, theta, alpha, and beta bands for wakefulness (j), NREM (k), and REM (l) stages in control and PLX5622-treated mice during the “light on” periods. **(m-o)** Normalized EEG power during wakefulness (m), NREM (n) and REM (o) in control and PLX5622 mice across the “light off” periods. **(p-r)** Quantification of EEG power in delta, theta, alpha, and beta bands for wakefulness (p), NREM (q) and REM (r) in control and PLX5622 mice during “light off” periods. n=7 mice for control group, and n=6 mice for PLX5622-treated group. Data are means ± SEM (bar graphs). Unpaired Student’s *t* test for **(a-f, j-l, p-r)**, \**p*<0.05, \*\**p*<0.01 and \*\*\**p*<0.001.

**Extended Data Fig 3.**
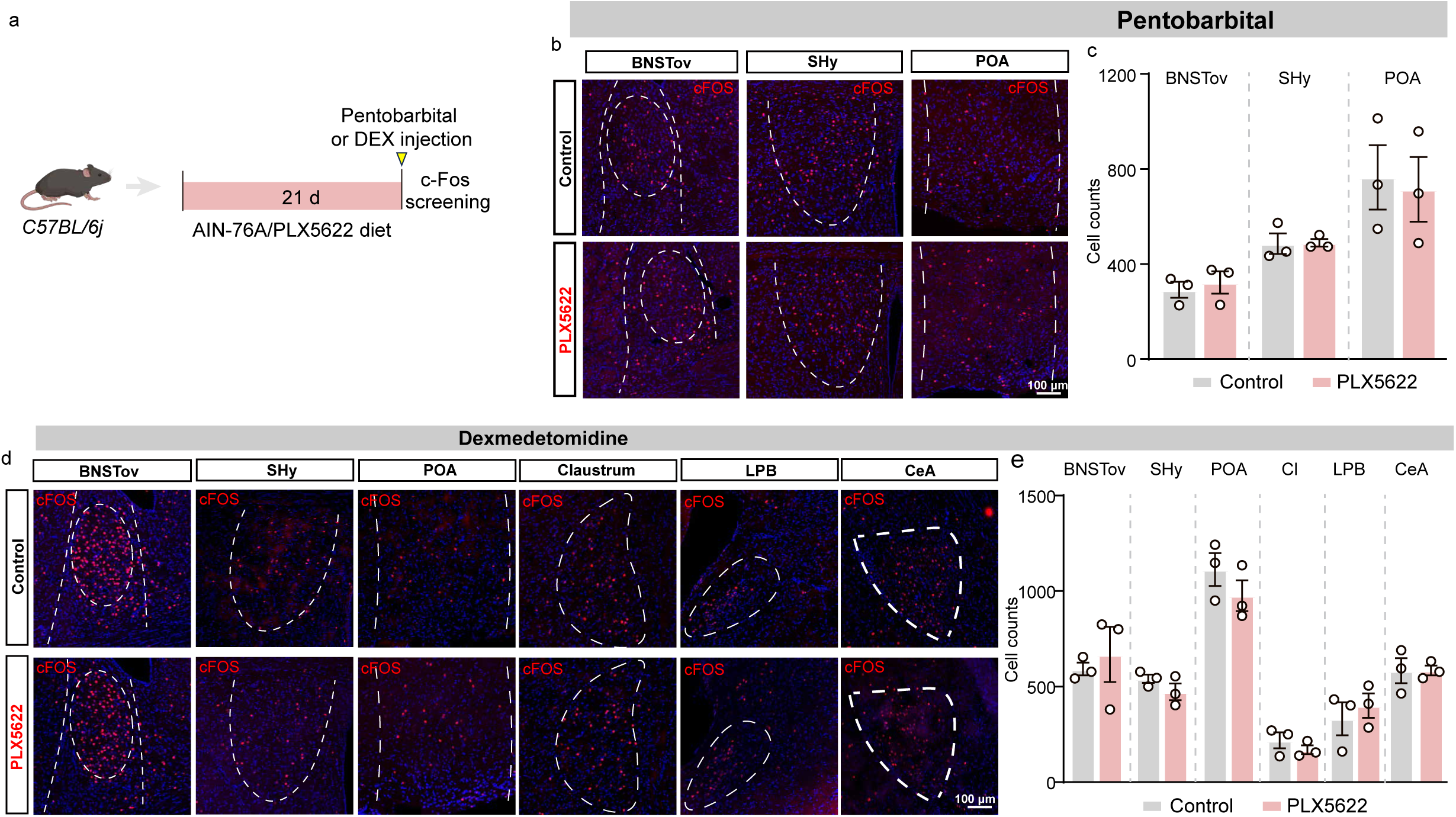
Microglial depletion did not affect the cFos staining pattern in the anesthetized brain. **(a)** A scheme of experimental design and the timing of PLX5622 treatment, anesthetics administration, and cFos screening. **(b)** Representative micrographs of cFos (red) in the BNSTov, SHy, and POA, in the control and PLX5622 group 1.5 hours following pentobarbital administration (100 mg/kg). **(c)** Number of cFos-positive neurons in selected anatomical sites after pentobarbital administration in the control and PLX5622 group. n=3 mice for control and PLX5622-treated group. **(d)** Representative micrographs of cFos (red) in the BNSTov, SHy, POA, Claustrum, LPB, and CeA in the control and PLX5622 group 1.5 hours following DEX administration (0.2 mg/kg). **(e)** Number of cFos-positive neurons in selected anatomical sites after DEX administration in the control and PLX5622 group. Data are means ± SEM (bar graphs). Unpaired Student’s *t* test for (**c**, **e)**, \**p*<0.05. BNSTov: bed nucleus of the stria terminalis, oval part; Shy: septohypothalamic nucleus; POA: preoptic area; LPB: lateral parabrachial nucleus; CeA: central amygdala.

**Extended Data Fig 4.**
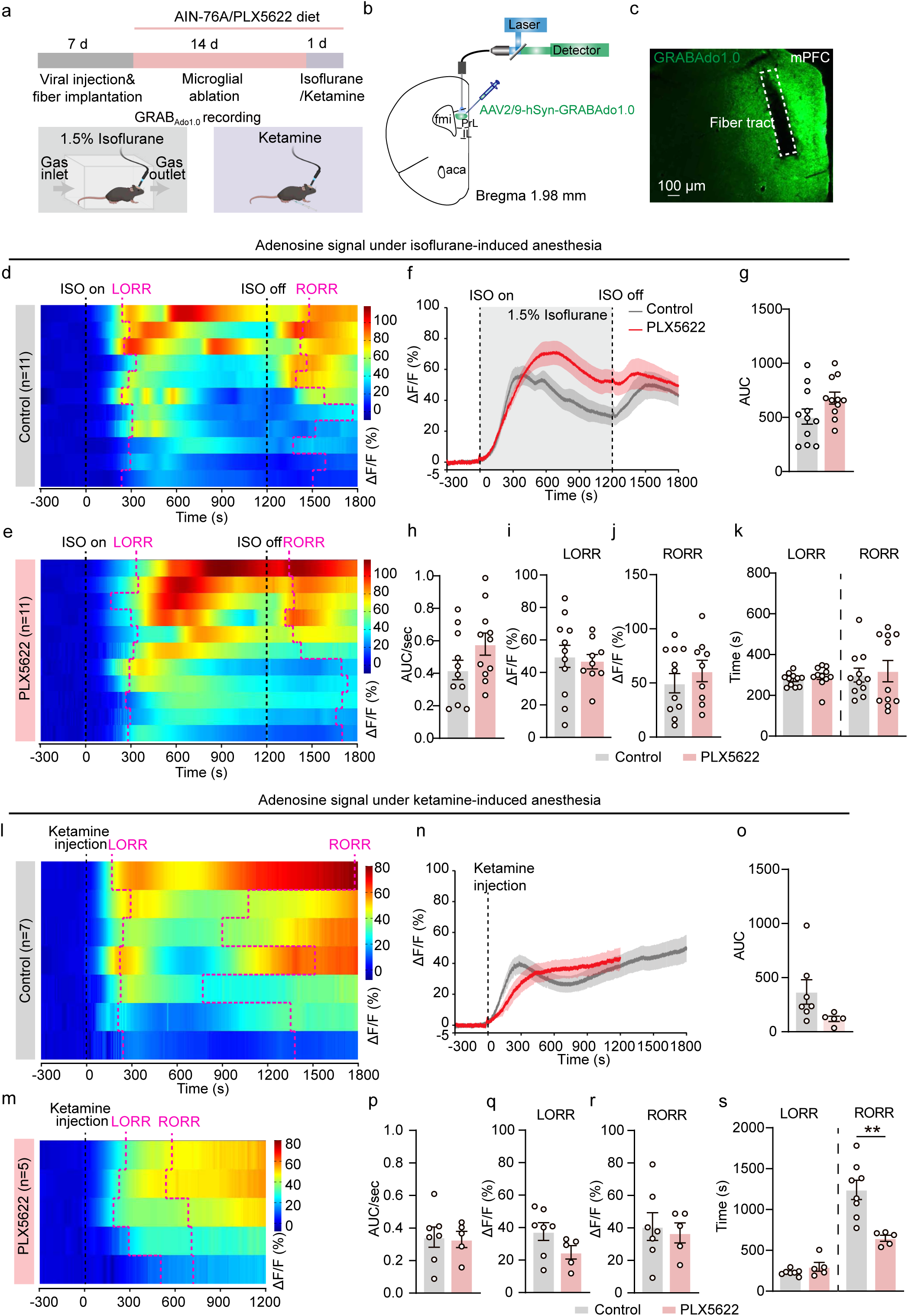
Extracellular adenosine levels barely change after microglial depletion during general anesthesia. **(a)** A schematic diagram depicting fiber photometry recording of extracellular adenosine levels during anesthesia. **(b)** Scheme of virus injection (adenosine sensors, GRAB_Ado_) and fiber photometry in the mPFC. **(c)** Representative sections showing the expression of adenosine sensors and fiber tract in the mPFC. **(d-e)** Heatmaps of extracellular adenosine signals during 1.5% isoflurane-induced anesthesia in control **(d)** and PLX5622-treated **(e)** mice. **(f)** Extracellular adenosine dynamics during isoflurane-induced anesthesia in the mPFC. **(g-h)** Quantification of AUC (g) and AUC/s **(h)** of adenosine dynamics between LORR and RORR during isoflurane-induced anesthesia. **(i-j)** Quantification of the adenosine levels at the time of LORR **(i)** and RORR **(j)** in isoflurane-anesthetized mice (n=11 mice). **(k)** Time of LORR and RORR after inhalation of 1% isoflurane in control and microglia-depleted (PLX5622-treated) male mice (n=11 mice per group). **(l-m)** Heatmaps of extracellular adenosine signals during ketamine-induced anesthesia in control **(l)** and PLX5622-treated **(m)** mice. **(n)** Extracellular adenosine dynamics during ketamine-induced anesthesia in the mPFC. **(o-p)** Quantification of AUC **(o)** and AUC/s **(p)** of adenosine dynamics between LORR and RORR during ketamine-induced anesthesia. **(q-r)** Quantification of the adenosine level at the time of LORR **(q)** and RORR in the ketamine-anesthetized mice **(r)** (n=7 for Control group; n=5 for PLX5622 group). **(s)** Time of LORR and RORR after injection with ketamine (50 mg/kg) in control and microglia-depleted (PLX5622-treated) male mice (n=7 for Control group; n=5 for PLX5622 group). Data are represented as mean ± SEM (bar graphs). Unpaired Student’s *t* test for **(g-k, o-s)**, \**p* < 0.05, \*\**p* < 0.01, and \*\*\**p* < 0.001. ISO: isoflurane; AUC: area under curve.

**Extended Data Fig 5.**
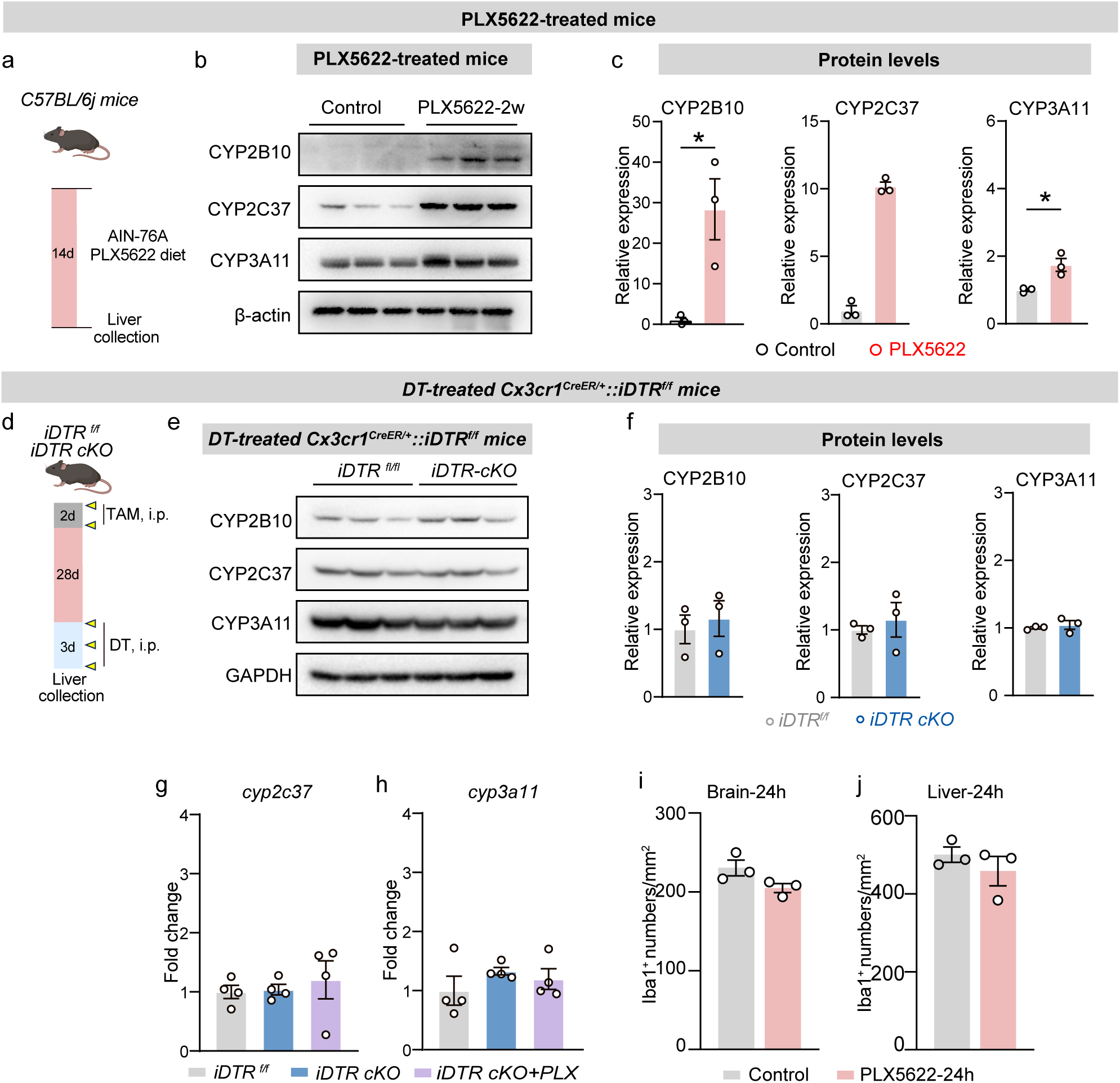
PLX5622 treatment induces the expression of CYP2B10 in the liver. **(a)** The experimental scheme and timing of PLX5622 treatment, and liver collection. **(b-c)** Representative blots (b) and quantification (c) of protein levels of CYP2B10, CYP2C37, and CYP3A11 in the liver from control and PLX5622-treated mice. n=3 mice per group. **(d)** The experimental scheme and timing of tamoxifen administration, DT treatment, and liver collection. **(e-f)** Representative blots (e) and quantification (f) of protein levels of CYP2B10, CYP2C37, and CYP3A11 in the liver from *iDTR^f/f^* and *iDTR cKO* mice. n=3 mice per group. **(g-h)** RT-qRCR analysis of *cyp2c37* (g) and *cyp3a11* (h) in the liver from *iDTR^f/f^*, *iDTR cKO*, and *iDTR cKO* with PLX5622-treated mice. n=4 mice per group. **(i-j)** Quantification of microglial number in the cerebral cortex (b) and liver (c) from control, and 24h PLX5622-treated mice. n=3 mice per group. Data are represented as mean ± SEM (bar graphs). Unpaired Student’s *t* test for (**c, f, i-j**), Mann-Whitney test for (**c, CYP2C37**), one-way ANOVA with Bonferron’s post hoc test for (**g-h**), \**p* < 0.05, \*\**p* < 0.01, and \*\*\**p* < 0.001.

**Extended Data Fig 6.**
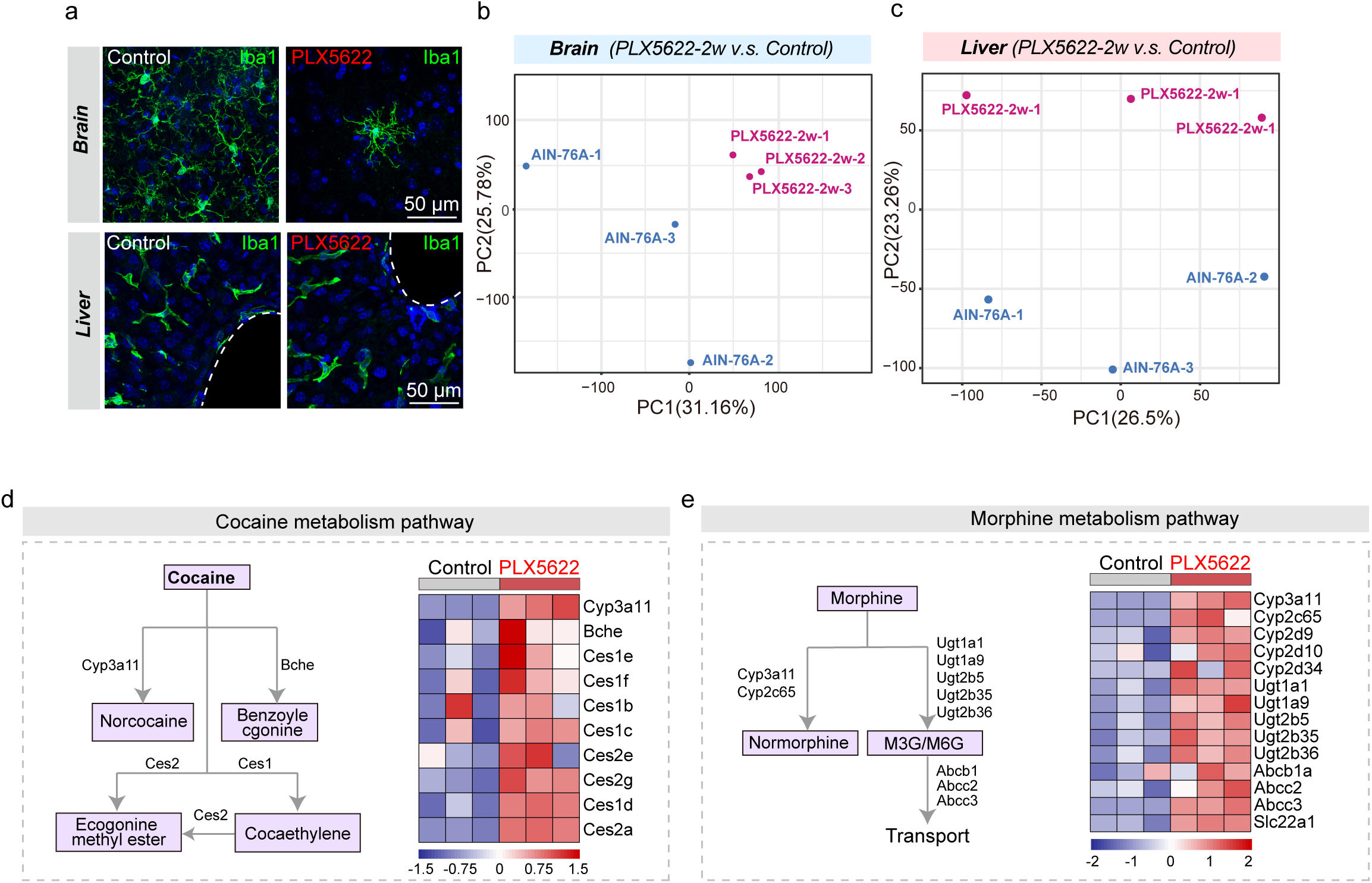
Bulk RNA sequencing of brain and liver tissue from control and PLX5622-treated mice. **(a)** Representative images showing the Iba1 staining from the brain and liver of the control and PLX5622-treated mice. **(b-c)** Principal component analyses showed the percentage of explained variance in the brain **(b)** or liver **(c)** between control and PLX5622-treated mice. **(d)** Heatmap of cocaine metabolism-pathway genes in the liver from the control and PLX5622-treated mice. **(e)** Heatmap of morphine metabolism-pathway genes in the liver from the control and PLX5622-treated mice.

**Table S1.**
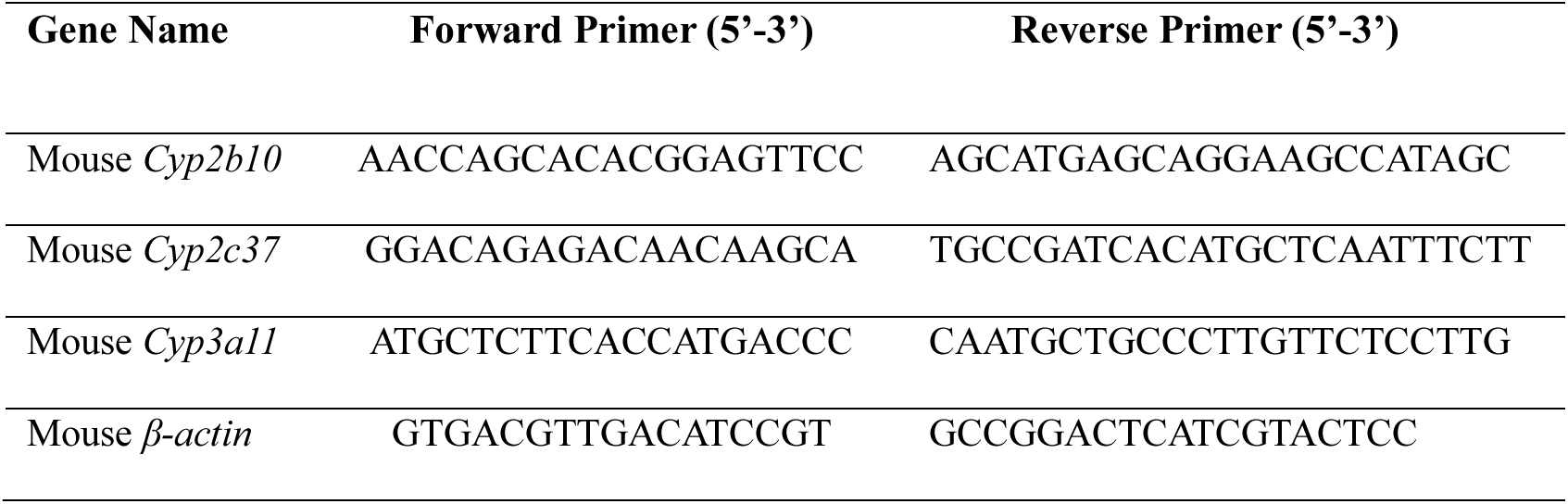
Primer sequences in RT-PCR.

**Table S2.** List of antibodies.

| **Antibody** | **Company/Origin** | **Catalog number** | **Application** | **Dilution** |
| --- | --- | --- | --- | --- |
| Rabbit anti-Iba1 | Wako | Cat # 019-19741 | IHC | 1:800 |
| Rabbit anti-cFOS | Synaptic Systems | Cat#226003 | IHC | 1:5000 |
| Mouse anti-β actin | Sigma-Aldrich | Cat # A1978 | WB | 1:1000 |
| Alexa Fluor 488-donkey anti-goat | Invitrogen | Cat # A11055 | IHC | 1:1000 |
| Alexa Fluor 555-donkey anti-rabbit | Invitrogen | Cat # A31572 | IHC | 1:1000 |
| Rabbit anti-CYP2B10 | Thermo Fisher | Cat#PA5-35032 | WB | 1:1000 |
| Rabbit anti-CYP2C37 | Abcam | Cat#Ab137015 | WB | 1:1000 |
| Rabbit anti-CYP3A11 | Thermo Fisher | Cat#PA1-343 | WB | 1:1000 |

